# Molecular basis of vitamin K-dependent protein γ-glutamyl carboxylation

**DOI:** 10.1101/2025.02.09.637231

**Authors:** Qihang Zhong, Dandan Chen, Jinkun Xu, Yao Li, Yan Meng, Qi Wen, Qiwei Ye, Guopeng Wang, Kexin Pan, Lin Tao, Jie Qiao, Jing Hang

**Affiliations:** State Key Laboratory of Female Fertility Promotion, Center for Reproductive Medicine, Department of Obstetrics and Gynecology, Peking University Third Hospital, Beijing, China; National Clinical Research Center for Obstetrics and Gynecology (Peking University Third Hospital), Key Laboratory of Assisted Reproduction, Ministry of Education (Peking University), Beijing key Laboratory of Reproductive Endocrinology and Assisted Reproduction, Beijing, China; Department of Biochemistry and Biophysics, School of Basic Medical Sciences, Peking University, Beijing, China; Department of Physiology and Pathophysiology, School of Basic Medical Sciences, Peking University, Beijing, China; Beijing Academy of Artificial Intelligence, Beijing, China; School of Life Sciences, Tsinghua University, Beijing, China; Department of Orthopedics, First Hospital of China Medical University, Shenyang 110001, China; State Key Laboratory of Membrane Biology, School of Life Sciences, Peking University, Beijing, China; Institute of Advanced Clinical Medicine, Peking University, Beijing, China

## Abstract

Gamma-glutamyl carboxylase (GGCX) in human is the sole enzyme responsible for catalyzing the carboxylation of glutamate (Glu) to γ-carboxyglutamate (Gla), a process that relies on the deprotonation of reduced vitamin K. This post-translational modification of vitamin K-dependent proteins (VKDPs) plays a pivotal role in numerous biological processes. Mutations in GGCX are responsible for multiple clinical phenotypes; however, the molecular mechanisms of GGCX activity and its pathology remain poorly understood. In this study, we reported the cryo-electron microscopy (cryo-EM) structures of GGCX bound with the either FIX or FX, two VKDPs and vitamin K, and employed the biochemical and cellular assays as well as molecular dynamics simulations to elucidate its dual-catalytic properties. The enzyme comprises a transmembrane domain that anchors the vitamin K-binding site, along with an ER luminal domain that facilitates binding of coagulation factor IX (FIX) at the exosite. The catalytic center orchestrates the oxidation of vitamin K and the carboxylation of Glu residues through a Cap-H2 coupling mechanism, along where disease-associated mutations in GGCX are predominantly clustered. These findings offer a comprehensive molecular framework for understanding the allosteric regulation of GGCX’s dual enzymatic activities and provide valuable insights for developing strategies to monitor and therapeutically target vitamin K metabolism in related diseases.

## Introduction

Gamma-glutamyl carboxylase (GGCX), also known as vitamin K-dependent (VKD) carboxylase, is a transmembrane enzyme embedded in the endoplasmic reticulum (ER)^1^. catalyzes the modification of glutamate(Glu) residues to γ-carboxyglutamate (Gla)^1^, a reaction coupled with the oxidation of vitamin K hydroquinone (VKH_2_) to its 2,3-epoxide form (VKO)^2, 3^. Mutations in *GGCX* or defects of VKD carboxylation are primary causes of bleeding disorders, collectively known as hereditary combined vitamin K-dependent coagulation factor deficiency (VKCFD)^4, 5^. VKCFD is an autosomal recessive disorder that is often life-threatening, with symptoms extending beyond bleeding to include developmental and skeletal anomalies. Beyond its role in the blood-clotting system, GGCX-mediated carboxylation is implicated in diverse biological processes, including vascular calcification, bone metabolism, cancer cell proliferation, and spermatogenesis^6–11^, and most recently the swine influenza virus adaption^12^, underscoring its significance as an interesting research target.

VKD proteins (VKDPs) must undergo carboxylation to achieve functional activity. Prominent VKDPs include procoagulants (e.g. coagulation factors VII, coagulation factors IX (FIX), and coagulation factors X (FX)), anticoagulants (e.g. proteins C, protein S, and protein Z), and proteins involved in bone formation (e.g. osteocalcin, also known as BGP) and soft tissue mineralization (e.g. matrix Gla protein, MGP)^13^. These proteins share a conserved N-terminal 18-residue propeptide that facilitates binding to GGCX^14^. The GLA domain, which contains the catalyzed glutamates, is located downstream of the propeptide. Interestingly, the propeptide binds at an exosite distinct from the catalytic site^15^. Early studies identified residues Arg234, Arg406, and Arg513 as key binding sites^16^, and subsequent research localized the binding site to the C-terminal region of GGCX, specifically between residues Pro495 and Arg513^17^. Despite these insights, the precise exosite regions and their coordination with catalytic activity remain incompletely understood.

As a dual-functional enzyme, GGCX simultaneously oxidizes vitamin K and carboxylates VKDPs. For its substrates, the glutamyl residues targeted for carboxylation are located downstream of the propeptide, within a region termed the GLA domain. Conversion of these Glu to Gla, followed by the cleavage of the propeptide, generates a calcium-binding module critical for the VKDP activation^18^. This reaction is initiated by the deprotonation of VKH_2_, enabling its reaction with oxygen to generate a carboxyl group required for subsequent carboxylation. Lys218 has been identified as the active base essential for the catalytic deprotonation of VKH ^19^. Furthermore, several disease-related mutations highlight the functional importance of key residues in GGCX^20^. For example, L394R mutation disrupts Glu binding at the active site^21^. Similarly, W501S impairs free propeptide binding^22, 23^, and W157R leads to a loss of vitamin K oxidation activity^24^. Nevertheless, the lack of structural data has long hindered a mechanistic understanding of the oxidation-carboxylation coupling mechanism and the pathogenic basis for GGCX mutations.

In this study, we present three high-resolution single-particle cryo-EM structures of wild-type (WT) and double-mutant (K217A & K218A) GGCX in complex with FIX or FX at 2.78, 2.59, and 2.58 Å resolution, respectively. By combining these structural data with *in vitro* and cell-based functional assays, as well as computational analyses, we elucidate the molecular basis of GGCX substrate binding and carboxylation activity. These findings not only shed light on the catalytic mechanism of GGCX but also elucidate the significant functions of key pathological mutations, highlighting the potential applications in advancing clinical anticoagulant therapy.

## Results

### Biochemical and structural analysis of the human GGCX•FIX complex

The full-length human GGCX protein was transiently expressed in HEK293F cells and purified using anti-FLAG affinity and size-exclusion chromatography (Supplementary Fig. 1a). The purified protein was then employed in an *in vitro* carboxylation reaction using FIX as the substrate (**Fig. 1a, b**). Consistent with previous studies^25, 26^, only vitamin K hydroquinone (VKH_2_) effectively promoted FIX carboxylation (**Fig. 1b**). Further concentration-gradient and time-lapse *in vitro* reactions validated the robust carboxylation activity of the purified GGCX protein (Supplementary Fig. 1b, c). To capture the structure of GGCX, we co-expressed GGCX with FIX to form a stable GGCX•FIX complex, which was subsequently purified to homogeneity in detergent (Supplementary Fig. 2a). Cryo-EM analysis of this complex yielded a structure at a resolution of 2.78 Å (**Fig. 1c**; Supplementary Fig. 2b–e; Table S1). The overall architecture of GGCX mainly consists of an N-terminal transmembrane domain (TMD; S32–N392) with nine transmembrane helices (TM1–TM9) and two luminal sub-domains oriented towards the ER lumen (**Fig. 1d, e**; Supplementary Fig. 3a). The luminal domain includes the propeptide-binding domain (PBD; S405–V604), comprising 11 β-sheets and two helices (H1 and H2), and the Arch domain (N605–A730), containing four helices (H3–H6) (**Fig. 1e**). H3 aligns parallel to the membrane, while H4 is perpendicular, and H5 lies on the detergent micelle surface (**Fig. 1d**). The cytosolic region is relatively short, comprising merely a few short α-helixes.

**Fig. 1.**
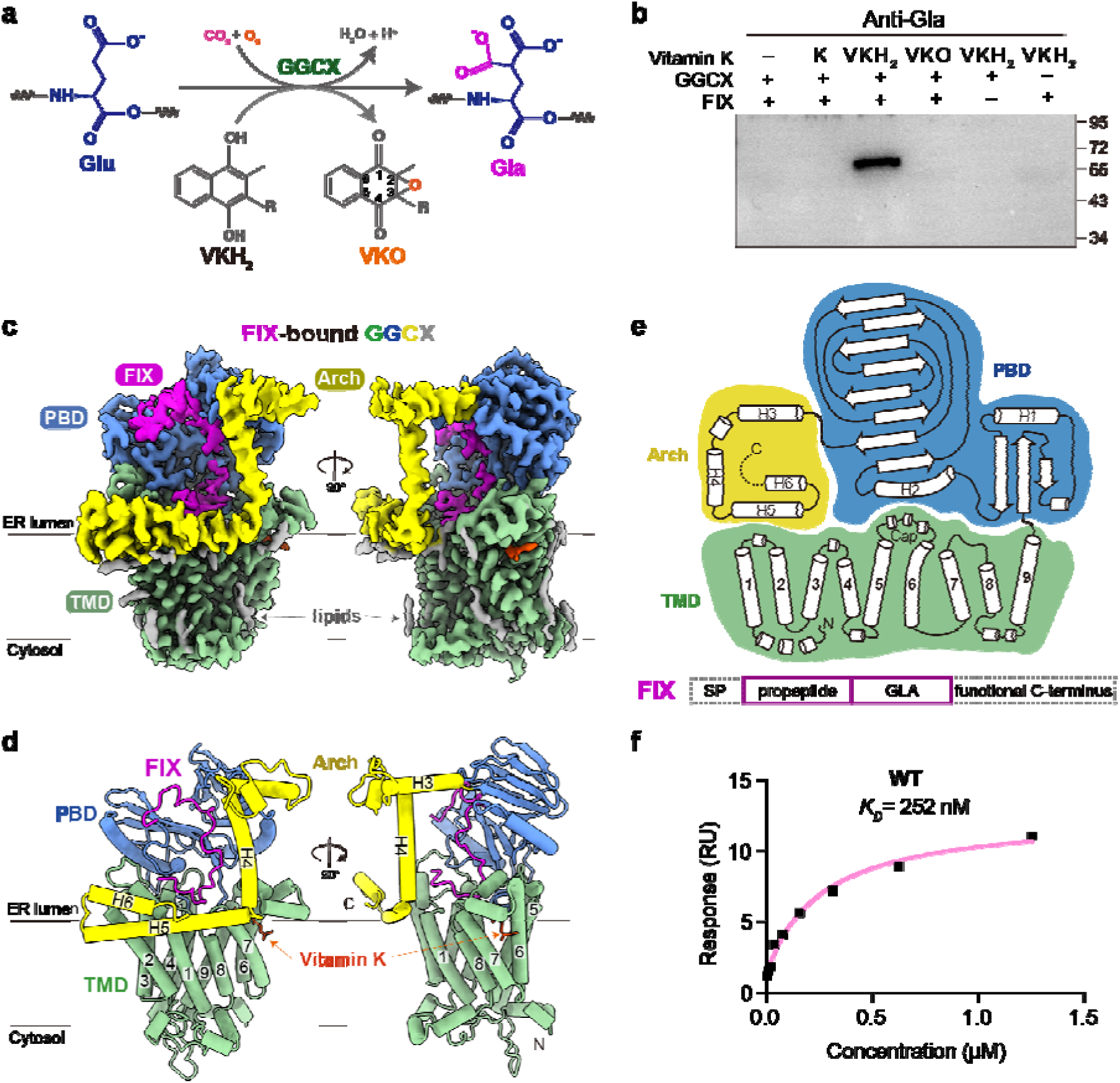
Overall structure of GGCX bound to Factor IX (FIX). **a,** Schematic representation of the carboxylation reaction catalyzed by GGCX, converting glutamte (Glu) to γ-carboxyglutamate (Gla). The reaction involves vitamin K hydroquinone (VKH_2_) as cofactor, which is oxidized to vitamin K epoxide (VKO). R, isoprenyl side-chain. **b,** The enzymatic activity assay of GGCX in the presence of FIX and different forms of vitamin K (K, VKH_2_, and VKO). VKH_2_ was generated by DTT-mediated reduction of menaquinone-4. The carboxylation products were detected using an anti-Gla antibody following electrophoresis. **c,** The cryo-EM map of the human GGCX bound to FIX and vitamin K is presented in a membrane view with a rotation of 90°. Individual domains are labeled and color-coded: the transmembrane domain (TMD) in green, the propeptide-binding domain (PBD) in blue, the Arch in yellow, coagulation factor IX (FIX) in purple, vitamin K in orange, and the surrounding lipids in gray. **d,** A cartoon representation of the FIX-bound GGCX structure in the same orientation as in **c**. The transmembrane segments and luminal helices are numbered and labeled. The color scheme is consistent across all figures. **e,** The topology map of human GGCX (top) and a schematic diagram of substrate FIX (bottom). The GGCX transmembrane segments are numbered (1–9), and the luminal helices are labeled (H1–H6). The structural elements of FIX are annotated. SP, signal peptide; GLA, carboxylated GLA domain. **f,** Surface plasmon resonance (SPR) analysis of GGCX binding to the propeptide and GLA domain of FIX (FIXQ/S). The dissociation constant (K*_D_*) was determined by analyzing the data with the steady-state affinity model using Biacore 8K+ Evaluation Software.

The cryo-EM density also allowed unambiguous assignment of the FIX substrate within the luminal domain cavity, specifically its propeptide and GLA domain (aa29–92) (**Fig. 1d, e**; Supplementary Fig. 3b). Surface plasmon resonance (SPR) analysis demonstrated strong binding between GGCX and the truncated FIX comprising the propeptide and GLA region (FIXQ/S), with a dissociation constant (*K*_D_) of 252 nM (**Fig. 1f**; Supplementary Fig. 1d). Meanwhile, an extra density located between TM5–TM8 and the elongated loop connecting TM5 and TM6 was discovered and identified as the endogenous vitamin K, which will be discussed further below (**Fig. 1c, d**). Numerous surrounding lipid densities and glycosylation sites were also discerned in the structure (**Fig. 1c**; Supplementary Fig. 3).

### ER Luminal domain is stabilized by interactions with TMD

Our structural analysis of GGCX reveals a distinctive feature within its ER luminal domain: the Arch-like domain (**Fig. 2a, b**). While the Arch motif is commonly associated with DNA or RNA helicase^27, 28^, its presence in membrane-embedded enzymes has been scarcely reported. Helix H5, an amphipathic structure within Arch, anchors to the membrane surface through its hydrophobic side which is embedded in the lipid bilayer (**Fig. 2a**). In addition to hydrophobic interactions, H5 also forms both direct and indirect interactions with the TMD. On one end of H5, a local density corresponding to a phospholipid was observed at the interface between H5 and TM7, with the lipid’s polar headgroup anchored by positive residues Arg680, Arg687, and Lys688 from H5 and forming a hydrogen bond with Ser291 of TM7 (**Fig. 2b, c**). On the opposite end, Arg694 is hydrogen-bonded with Asp89 of TM1, Ser702 and Asn705 form hydrogen bond networks with Asp113 of TM2 (**Fig. 2c**). These interactions tether H5 to the TMD, constraining the swing movement of Arch. To explore the roles of Arch and the invisible C-terminal region, two GGCX variants were generated: a C-terminal region deletion (ΔC) or a CTD&H5&H6-deletion (ΔArch) construct. *In vitro* carboxylation assays demonstrated that ΔC retained most of its activity, showing only a mild reduction compared to the full-length GGCX (WT), whereas ΔArch exhibited a significant loss of catalytic function (**Fig. 2d**). Similar results were obtained from cell-based assay (**Fig. 2e**), indicating that while the relatively less conserved C-terminus may act as a regulatory element for FIX carboxylation, the conserved Arch is crucial for GGCX luminal domain assembly and the subsequent enzymatic activity (Supplementary Fig. 4).

**Fig. 2.**
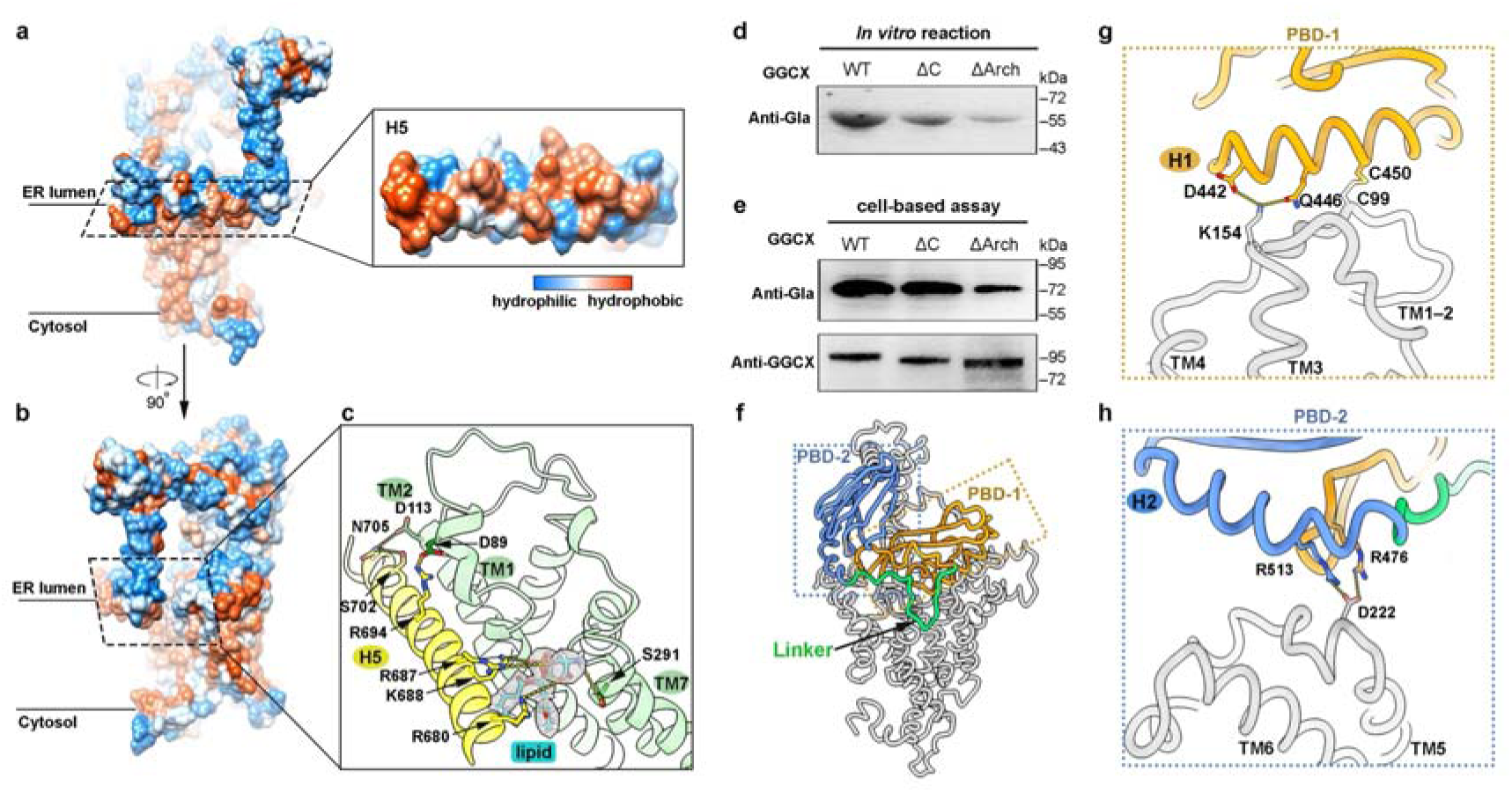
The luminal domain anchors to the TMD and the ER luminal leaflet. **a,** Surface hydrophobicity map of GGCX with the surface of H5 facing the membrane is magnified for clarity. **b,** A 90° rotation of the view shown in **a**. **c,** The interactions between helix H5 and the TMD are highlighted. The density of the lipid molecule that trapped between the H5 and TM7 is shown as gray transparent surface and cyan sticks. Residues involved in the lipid interactions on one end and residues mediating interactions between H5 and TM1–2 on the other end are labelled. Hydrogen-bonds are shown as yellow dashed lines. **d–e,** Carboxylation activity assays of truncated GGCX constructs, with either a C-terminal deletion (ΔC) or the CTD&H5&H6-deletion (ΔArch). Both in vitro reaction (**d**) and cell-based assay (**e**) were performed. **f,** Structural representation of GGCX in complex with FIX. The PBD consists of two subdomains: PBD-1 (residues 405–489) and PBD-2 (residues 497–604). PBD-1, PBD-2, and the linker between them are colored in orange, blue, and green, respectively. **g–h,** Molecular interactions between PBD-1 (**g**) and PBD-2 (**h**) with the TMD. Residues involved in their interactions are labelled and represented as sticks. The hydrophilic interactions are indicated as yellow dashed lines.

As the main functional luminal sub-domain, the PBD also interacts extensively with the TMD (**Fig. 2f–h**). PBD is divided into two lobes: PBD-1 (S405–V489) and PBD-2 (D508–V604) (**Fig. 2f**). PBD-1 is composed of three β-sheets and helix H1, with Cys450 of H1 forming a disulfide bond with Cys99 from the TM1–2 loop. Additionally, Asp442 and Gln446 of H1 form hydrogen bonds with Lys154 (**Fig. 2g**). PBD-2 consists of eight β-sheets and helix H2, where Arg476 from PBD-1 and Arg513 from H2 form salt bridges with Asp222 simultaneously (**Fig. 2h**), consistent with reports that Arg476 is critical for carboxylation activity with mutations leading to disease^20, 29^. The two PBD subdomains are connected by a flexible linker (Q490–M507), therefore forming a groove on the opposite face, which provides an ideal location for the following substrate binding (**Fig. 2f**).

Although these interactions contribute to stabilize the PBD and the Arch with the TMD, the intrinsic flexibility of the Arch potentially allows extensive luminal domain movement. To further investigate this hypothesis, we performed molecular dynamics (MD) simulations of apo GGCX, showing that the Arch helices rapidly collapsed inward toward the interior of the complex. This movement was coupled with a discernible propensity of vitamin K to dissociate from the binding pocket (Supplementary Fig. 5a; Supplementary Video 1). To quantify these structural dynamics, we calculated the per-residue root-mean-square-fluctuation (RMSF) (Supplementary Fig. 5b). The results showed that most residues of GGCX exhibited negligible fluctuations, with RMSF values of approximately 1 Å. In contrast, the C-terminal helices displayed significant deviations, exceeding 5 Å in all simulations, leading to an overall root-mean-square-deviation (RMSD) of approximately 8 Å (Supplementary Fig. 5c). These results underscore the importance of substrate binding for the structural stability of GGCX, highlighting that the enzyme undergoes dynamic conformational changes in the absence of its substrate. This flexibility likely enables the PBD to coordinate with various substrate interactions, and upon substrate binding, the conformation stabilizes, preparing GGCX for its catalytic activity.

### Exosite binding of propeptide by PBD

Next, we analyzed the binding properties between the propeptide and PBD. As mentioned above, PBD-1 and PBD-2 together form a claw-like structure that clamps the substrate within the central groove (**Fig. 3a**). Interestingly, this binding site is distinct from the active catalytic center where vitamin K binds and is therefore designated as an exosite (**Fig. 1d**). Within this binding interface, multiple hydrophilic interactions occur between the main chains of GGCX and the propeptide (**Fig. 3b**). Additionally, His34 of FIX engages in a π-π stacking interaction with Tyr425 and the benzene ring of Phe31 inserts into a hydrophobic pocket in BSD-1. Meanwhile, Leu41 establishes hydrophobical interactions with BSD-2, further stabilizing the binding (**Fig. 3b**). Sequence alignment of different substrates reveals the high conservation of Phe31 but more variability of His34 (Supplementary Fig. 6a), although the carboxylation activities of GGCX towards different substrates remain undistinguishable (Supplementary Fig. 6b, c). To evaluate the functional importance of these residues, we performed the site-directed mutagenesis and discovered that F31G, H34R and L41W mutants exhibited a significant reduction in the *in vitro* binding affinity (*K_D_* of ∼3–6 μM), over ten-fold lower than that of WT-GGCX (**Fig. 3c**; **Fig. 1f**; Supplementary Fig. 6d). Meanwhile, pull-down assay utilizing immobilized GGCX and GFP-tagged mutant FIX further validated these findings, as all tested mutations disrupted the propeptide’s binding to GGCX (**Fig. 3d**).

**Fig. 3.**
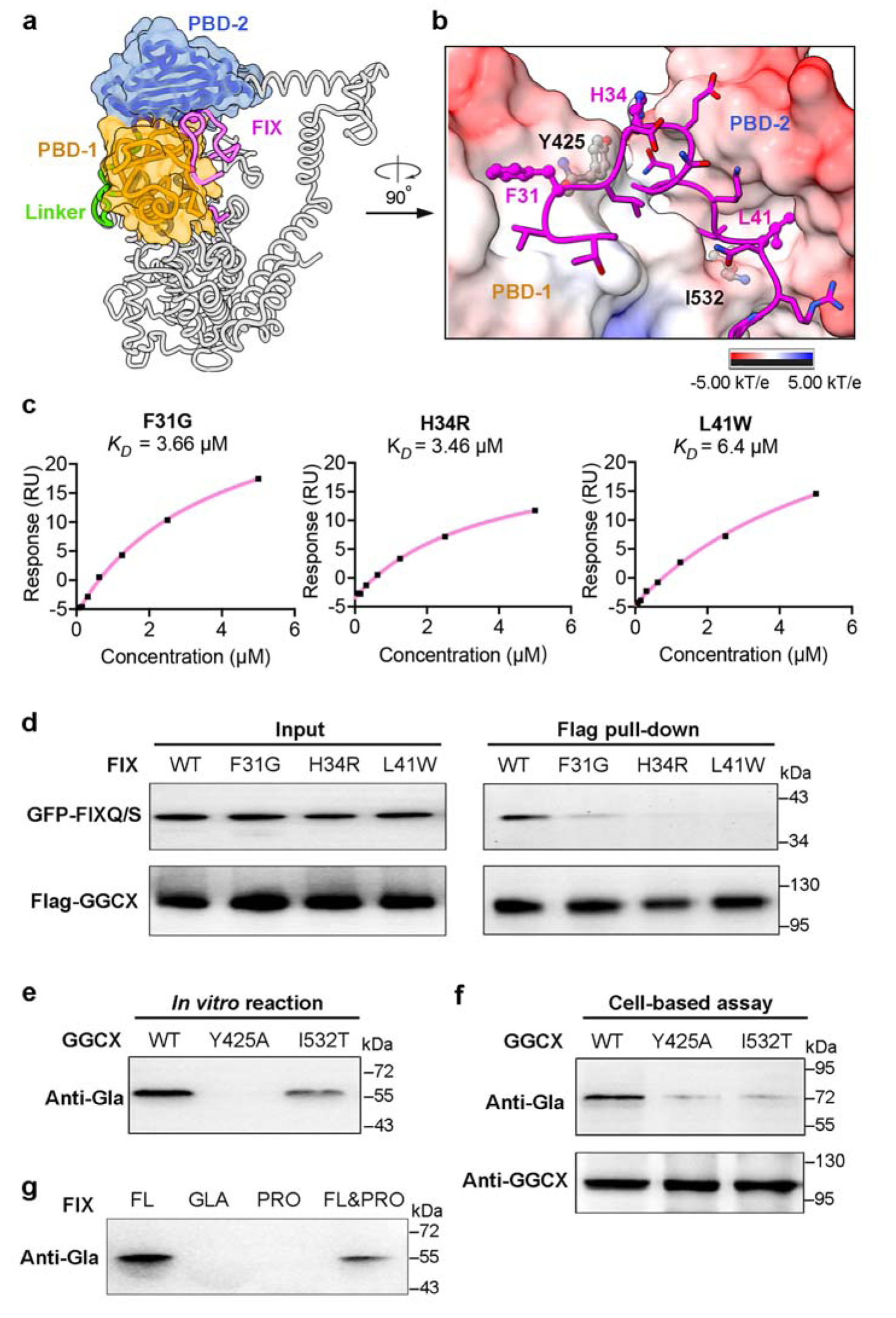
Critical molecular interactions between GGCX and the propeptide region of FIX. **a,** The propeptide of FIX is clamped inside the groove formed by PBD-1 and PBD-2. The surface of PBD-1 and PBD-2 is shown in orange and blue, respectively, with the linker between them colored in green. The bound FIX is colored in purple. **b,** The detailed interactions between the propeptide and PBD, with PBD displayed as surface charges. Key interacting residues in both GGCX and FIX are labeled and shown as ball-and-sticks. **c,** SPR-determined binding affinities between mutated FIX (F31G, H34R, and L41W) and GGCX. **d,** Pull-down assays for Flag-immobilized GGCX and GFP-fused mutated FIXQ/S. GFP signals were detected by fluorescence. **e–f,** Carboxylation activity of FIX using GGCX with loss-of-function mutations measured by both *in vitro* reaction (**e**) and cell-based assay (**f**). **g,** Carboxylation activity of FIX with different truncations. FL, full-length FIX; GLA, GLA domain of FIX; PRO, the propeptide region of FIX; FL&PRO, addition of excess free propeptide into the full-length FIX reaction.

To further investigate the role of specific residues in the interaction between GGCX and the propeptide region of FIX, we focused on two hydrophobic residues, Tyr425 and Ile532 (**Fig. 3b**). Mutational analysis revealed that the Y425A mutant showed a pronounced reduction in carboxylation activity, as demonstrated by both *in vitro* reaction and cell-based assays. In contrast, the I532T mutation exhibited a milder but still detectable effect on carboxylation activity (**Fig. 3e, f**). These findings indicate that I532T mutation disrupts substrate binding, thereby compromising GGCX function. This is the first mechanistic explanation for the pathologic basis of I532T, which has been associated with impaired hemostasis^30^. To further investigate the function of the propeptide, we performed a competition assay (**Fig. 3g**; Supplementary Fig. 6e). FIX containing only the GLA domain (GLA) or propeptide region (PRO) failed to undergo carboxylation (**Fig. 3g**); in the presence of excess free propeptide peptides (FL&PRO), carboxylation efficacy was significantly reduced, indicating that PRO competes with full-length FIX for binding to GGCX. Collectively, these results collectively demonstrate that the propeptide of FIX enhances GGCX-mediated carboxylation by facilitating substrate binding.

### Catalytic reaction center: integration of vitamin K and GLA-binding pockets

To facilitate its dual function, the catalytic reaction center of GGCX must simultaneously accommodate both the vitamin K-binding and GLA-binding. Cryo-EM analysis revealed that the GLA domain binds within an internal cavity formed by the luminal domain and the TMD (**Fig. 4a**). A cutaway surface view further illustrates that the pockets for vitamin K and GLA are interconnected, collectively forming what we designate as the catalytic reaction center (**Fig. 4b**). Intriguingly, our structural analysis also uncovered a distinct density embedded within an elongated pocket in the TMD, located in close proximity to Glu53 of FIX (**Fig. 4a–b**). Based on these observations and the density interpretation, we identified this molecule as vitamin K. However, while we could not definitively determine its precise form, we assigned it as (menaquinone-4, MK-4) in our model (Fig. 4b). To address this uncertainty, we designed a double mutant (K217A/K218A) of GGCX to abolish its enzymatic activity and subjected the protein to cryo-EM analysis. The structures of GGCX-K217A/K218A (GGCX^AA^) in complex with FIX and FX were determined at 2.59 Å and 2.58 Å, respectively (Supplementary Fig. 7; Supplementary Table 1). Both structures revealed a more defined EM map for vitamin K with high local resolution, enabling us to confidently assign it as the MK-4 hydroquinone form (MKH_2_-4) (**Fig. 4c–d**). To further clarify the molecular interactions, we superimposed the structure of GGCX•FIX onto the GGCX^AA^•FIX and modeled an MKH_2_-4 molecule into the GGCX•FIX complex (**Fig. 4e**). This refined model served as the basis for our detailed investigation hereafter.

**Fig. 4.**
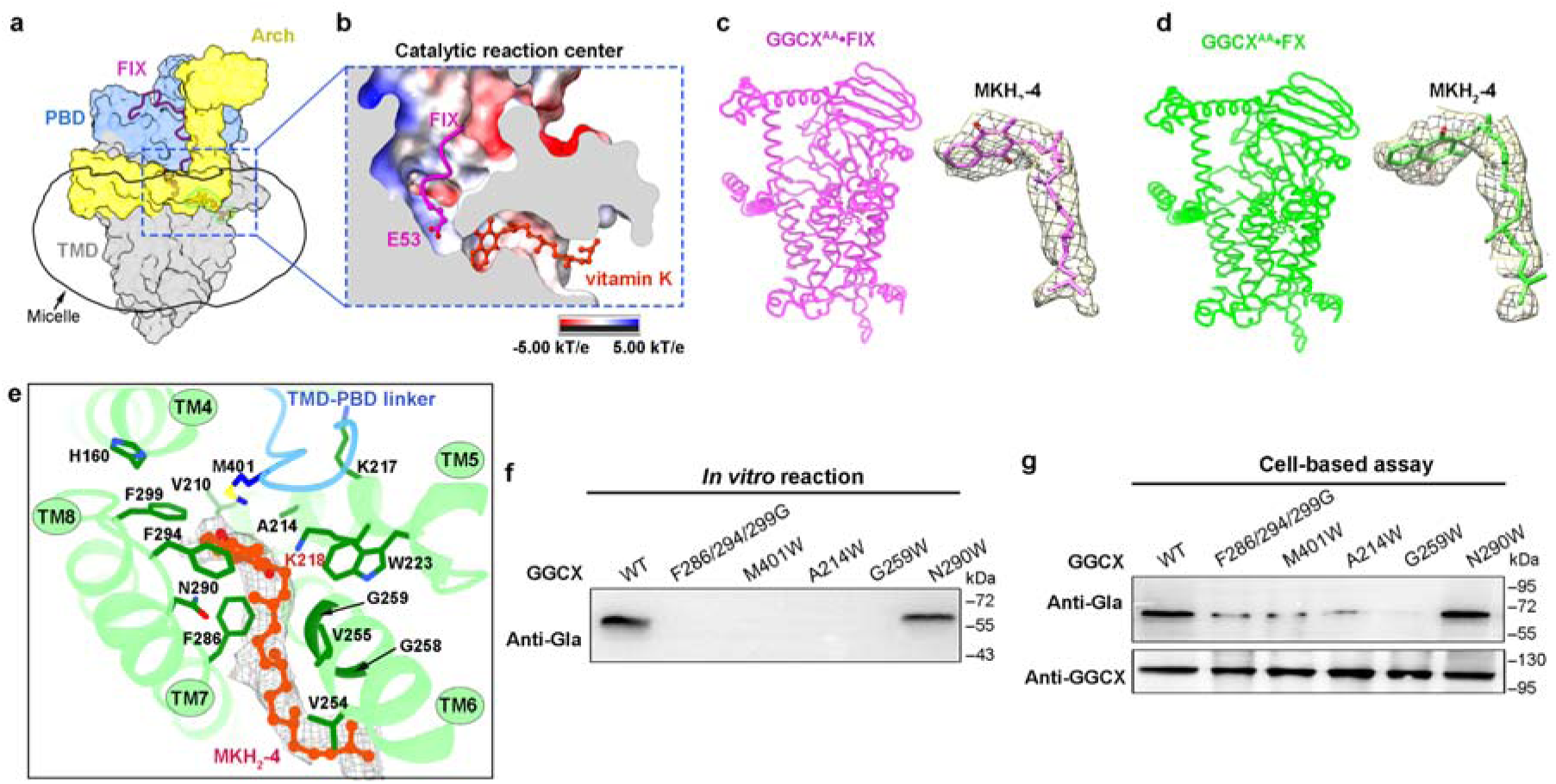
The vitamin-K binding site. **a,** Overall view of GGCX, FIX, and vitamin K. The surface of the TMD, PBD, and Arch domains are shown in gray, blue, and yellow, respectively. Vitamin K and substrate FIX are shown as cartoons and colored orange and purple, respectively. The detergent micelle is shown as a transparent surface indicated by an arrow. The reaction center is highlighted by a blue dashed box. **b,** Cutaway side-view of the reaction center. GGCX is displayed as electrostatic potential surface with positively-, negative-charged, and neutral residues colored in blue, red, and white, respectively. The bound substrate FIX is shown as a cartoon, and vitamin K is represented as a ball-and-stick model. **c–d,** Cryo-EM structures of K217A/K218A double mutant GGCX (GGCX^AA^) in complex with either FIX (GGCX^AA^•FIX complex) (**c**) or FX (GGCX^AA^•FX complex) (**d**). The densities for the bound vitamin K in MKH_2_-4 form are displayed. **e,** Detailed interactions between vitamin K and key residues. The density of vitamin K is shown as a mesh. Residues from the TMD are colored green, and the residue in the TMD-PBD linker is colored blue. **f–g,** Carboxylation activity of GGCX with key residue mutations assessed by *in vitro* reaction (**f**) and cell-based assay (**g**).

The orientation of MKH_2_-4 is parallel to the membrane, with its 2-methyl-1,4-naphthoquinone ring buried inside a hydrophobic tunnel, while its isoprenyl side chain extends toward the periphery (**Fig. 4b**). On the head side, the naphthoquinone ring is coordinated by a large hydrophobic pocket involving Phe286 from TM7, Phe294 and Phe299 from TM8, Val210 and Trp223 from TM5, His160 from TM4, and Met401 from TMD-PBD linker (**Fig. 4e**). On the tail side, the isoprenyl side-chain traverses a narrow tunnel (∼11 Å in diameter), which is lined by small residues such as Ala214 from TM5, Gly258 and Gly259 from TM6, Asn290 from TM7, and Val254 and Val255 from TM6 (**Fig. 4b, e**). To validate this vitamin K binding site and the role of these surrounding residues, we either deleted the side chains in the hydrophobic pocket (F286/F294/F299G), or introduced steric hindrance to the naphthoquinone ring (M401W), or inserted bulky residues into the isoprenyl tunnel (A214W, G259W, and N290W). These mutations significantly impaired carboxylation activity in both *in vitro* assays and cell-based experiments, except for N290W that showed partial activity (**Fig. 4f, g**). Beyond the reduction in activity, these loss-of-function mutations also resulted in a nearly 10-fold reduction in binding affinity for vitamin K compared to the WT protein, which has a *K_D_* of 0.85 μM (Supplementary Fig. 7). These findings provide compelling evidence that these mutations disrupt the integrity of the vitamin K-binding pocket, thereby confirming their critical role in creating a hydrophobic pocket that accommodated the vitamin K binding.

For GLA binding, the γ-carboxyl group of Glu53 from FIX is inserted into a positively charged binding pocket near Lys218, situated in close proximity to the naphthoquinone ring of vitamin K (**Figs. 4c, 5a**). This residue has been identified as the catalytic site by serving as the weak base to facilitate the deprotonation of VKH_2_ during reaction^19^. Structural analysis showed that Lys218 forms a hydrogen bond with the naphthoquinone of MKH_2_-4, reinforcing its role in catalysis (**Fig. 5b**). Additionally, Asn159, His160, and Tyr395 engage in hydrogen bonding with the γ-carboxyl group of Glu53, while Asn158 and Arg436 stabilize the main chain of the GLA domain, collectively constituting the Glu pocket (**Fig. 5b**). Previous studies have shown that the H160A mutation impairs Glu carboxylation^31^, underscoring the role of His160 in stabilizing the transition state during carbanion formation. In the meantime, Phe294, Phe299, Met401, and Met401 contribute to a “hole-in-the-wall”-like structure that separates the Glu pocket from the vitamin K pocket, positioning them on opposite sides (**Fig. 5b**). Since mature FIX adopts a well-folded structure after carboxylation^32^, the GLA domain observed in our cryo-EM structures likely represents an immature state.

**Fig. 5.**
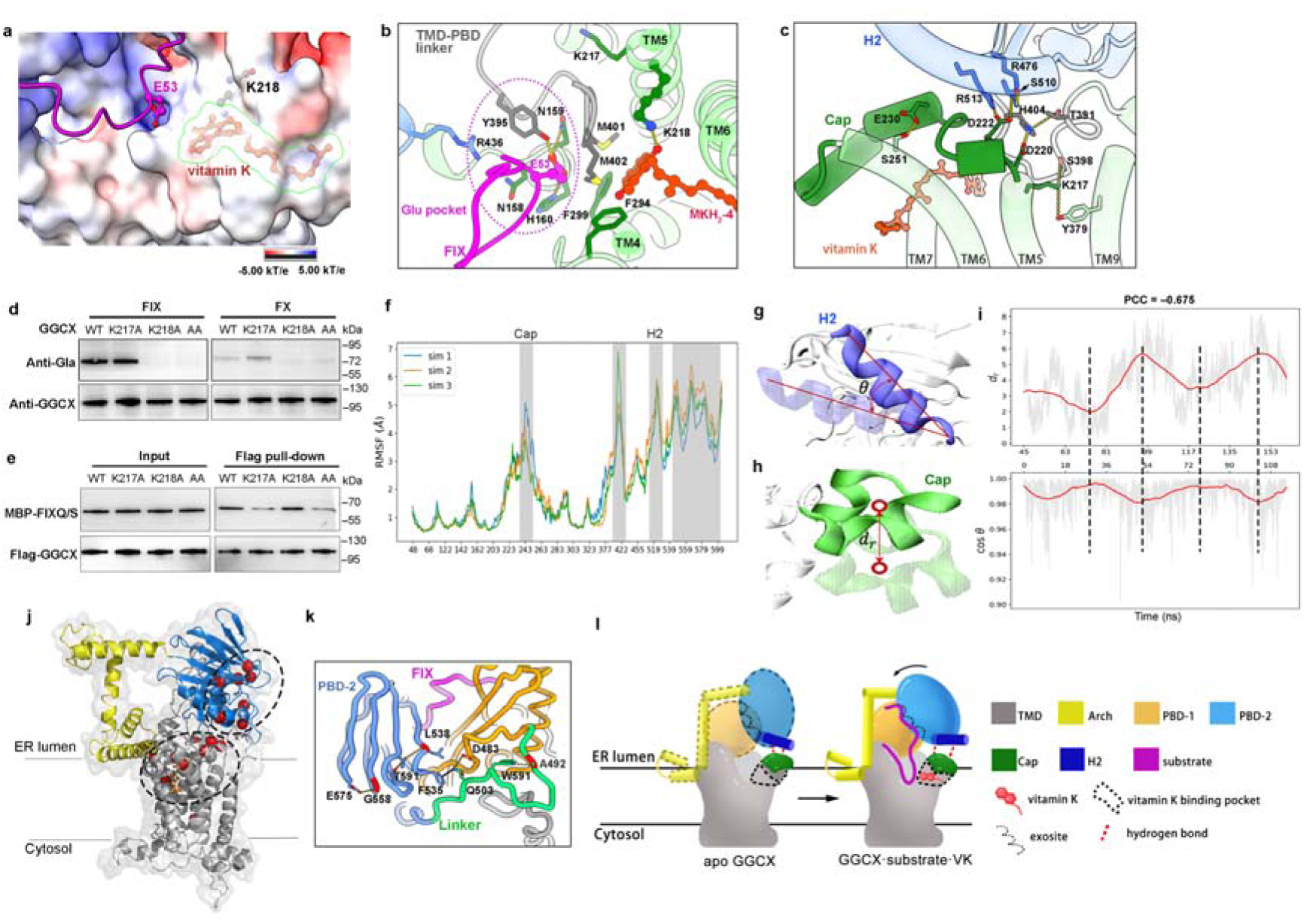
The reaction center of GGCX. **a,** The GLA region of FIX inserts into a positively charged pocket of GGCX near the vitamin K-binding pocket. GGCX is displayed as an electrostatic potential surface. E53 of FIX, K218 of GGCX, and vitamin K are shown as ball-and-sticks. The vitamin K-binding pocket is outlined with green line. **b,** The active reaction center of GGCX bound to FIX and vitamin K. The TM helices of GGCX are numbered and colored green, while the TMD-PBD linker is shown in gray. The Glu pocket is indicated by a dashed magenta oval. **c,** The interactions between the Cap and PBD-2. Cap region (D220 to S246) of GGCX are colored dark green. Residues involved in interactions between the TMD and H2 of PBD-2 (colored blue) are labeled. **d,** Cell-based carboxylation activity of different GGCX mutants towards either FIX or FX. **e,** Pull-down assays for Flag-immobilized GGCX with MBP-fused FIXQ/S region. **f,** Root mean square fluctuation (RMSF) of GGCX, excluding the C-terminal residues and Arch, were obtained from three independent accelerated molecular dynamics (aMD) simulations. The Cap and H2 are highlighted in light gray. **g–h,** A schematic representation of quantitative metrics. The angle (cos*θ*) to evaluate the motion of H2 (**g**), and the distance (*dr*) is used to quantify Cap movement (**h**). The initial reference position is rendered in transparency. **i,** Temporal evolution of *dr* (top) and cos*θ* (bottom) during the simulation. The original data are shown as grey transparent lines, while the red solid lines denote the smoothed trajectories. The two metrics are aligned after compensating for temporal delay, with peaks and valleys marked by black dashed lines. **j,** Diseases-associated mutations of GGCX are mapped onto the structure and highlighted as red spheres. **k**, Mutations locates within PBD form several hydrogen-bonds. Hydrogen-bonds and cation-π interaction are shown as yellow and black dashed lines, respectively. **l**, A working model of GGCX. In the apo form of GGCX, the luminal domain that exhibits significant conformational flexibility; substrate binding at the exosite and vitamin K binding is coordinated by the synergistic H2-Cap transduction mechanism.

Adjacent to Lys218 is another positively charged residue Lys217, which is regarded to modulate the activity of Lys218 and stabilize the carbanion group of the substrate’s glutamate^15^.Suprisingly, the cryo-EM structure shows that the ε-amino group of Lys217 is stabilized by hydrogen bonds with Ser398 and Tyr379 from TM9, positioning it approximately 12 Å away from the Glu’s carbanion group (**Fig 5c**). Given that the TMD-PBD linker (T391–H404), which locates between TM4 and TM9, is embedded within the membrane (**Fig 5b**), it is unlikely for Lys217 to form direct interactions with the glutamate side chain. Therefore, the primary function of Lys217 appears to be stabilizing the interaction between the TMD and PBD-2, thereby maintaining the structure integrity of the catalytically active enzyme. To further elucidate the function of these two key residues, we generated two single-point mutations. The K218A mutation resulted in a significant loss of activity both for FIX and FX (**Fig. 5d**), while its binding affinity for FIX remained unaffected (**Fig. 5e**). In contrast, the K217A mutation impaired FIX and FX binding but had no effect on catalytic activity (**Fig. 5d, e**). These results demonstrate that Lys218 is critical for the catalytic reaction, whereas Lys217 plays a key role in mediating conformational coupling.Importantly, this function is independent of the catalytic role of Lys218, highlighting the distinct roles of these two residues in regulation of GGCX activity.

### Coupling mechanism underlying the propeptide and vitamin K binding

In the surrounding area of vitamin K binding, the loop between TM5 and TM6 forms a Cap-like structure that encloses the vitamin K binding site, separating it from the ER luminal solvent (**Fig. 5c**). Structural analysis reveals an extensive hydrogen-bond network stabilizing this Cap, which interacts with H2 of PBD-2, involving residues such as Arg513-Asp222-Arg476-Ser510, Asp220-His404-Thr391, Glu230-Ser251, and Tyr379-Lys217-Ser398 (**Fig. 5c**). Based on these observations, we hypothesize that the binding or release of vitamin K influenced by the Cap is coupled with the movement of H2. To explore this hypothesis, we performed MD simulations using a GGCXΔArch-FIX-MK-4 system. Conventional MD simulation revealed little change in all repeats (Supplementary Fig. 9), so we employed accelerated MD (aMD) simulations to enhance conformational sampling. The system maintained overall structural rigidity, with the transmembrane helices remaining nearly rigid in majority of TMD (RMSF < 1 Å) (**Fig. 5d**; Supplementary Video 2). Remarkable conformational transitions were observed in both the H2 and Cap (**Fig. 5f–h**). To quantify these movements, we proposed novel metrics in the simulation trajectories: 1) the cosine of the angle between the H2 and its initial orientation (cos*θ*) (**Fig. 5g**), and 2) the deviation of the Cap’s center of mass from its initial position (*dr*) (**Fig. 5h**). In a 160 ns aMD simulation, the H2 underwent intermittent opening events (**Fig. 5g, i**), and the Cap exhibited intermittent conformational transitions (**Fig. 5h, i**). Notably, the motions of the H2 and Cap were not synchronized, with a temporal delay of around 45 between the two processes. Once the temporal delay was removed, these two processes showed strong correlation, with a Pearson correlation coefficient (PCC) of –0.675 (**Fig. 5g**). Furthermore, Superimposition of the three cryo-EM structures showed that while the TMD remained unchanged, the PBD-2 in the GGCX^AA^•FIX and GGCX^AA^•FX complex adopted a ∼5°-rotation away from the active center compared to GGCX^WT^•FIX structure (Supplementary Fig. 10a). This rotation is likely due to the loss of Lys217, which disrupts the connection between the Cap and PBD-2. These results suggest that H2 acts as a conduction element coupled with the conformational transition of Cap, with the lagged yet synergistic interaction between these two elements establishing a specific H2-Cap transduction mechanism.

As aforementioned, missense mutations in GGCX are associated with various clinical disease phenotypes^33^ (Supplementary Table 2). Mapping their locations onto the GGCX structure disclosed two main clusters: one at the catalytic center and the other at the interface between PBD-1 and PBD-2 (**Fig. 5j**). Key residues in this interface, including Trp501, Thr591, and Gly558, are mainly involved in forming hydrogen-bond networks within the PBD (**Fig. 5k**). Together, these results suggest that propeptide-binding by PBD is linked to the movement of Cap that influences vitamin K binding, facilitating the coupling between the catalytic site and propeptide binding.

## Discussion

Vitamin K serves as a cofactor by the enzyme GGCX in the synthesis of Gla proteins through a process known as carboxylation. Carboxylated proteins have the ability to chelate calcium ions, which are essential for blood coagulation and for controlling binding of calcium in bones and other tissues^34^. Deficiency of vitamin K can lead to serious consequences such as VKCFD, osteoporosis, and calcification of arteries^35^. Although many previous genetic studies have identified key residues related to GGCX activity, no mechanistic illustration exists to date. Remarkably, GGCX exhibits no known homology to any other enzyme family, highlighting its unique structural and functional characteristics. Through structural determination and direct experimental evidence, our study uncovers the molecular basis underlying Gla carboxylation as well as vitamin K epoxidation. The coupling of two half-reactions in a 1:1 stoichiometry remains a fundamental concept in understanding the enzymatic activity of GGCX (**Fig. 5l**).

One fascinating aspect of this study is the exploration of the exosite-binding of the propeptide. Our attempts to obtain the apo-GGCX structure were unsuccessful (Supplementary Fig. 5a), which implies that the binding of the propeptide enhances the stability of GGCX luminal domain, in line with previous studies^36^. This mechanism is achieved through allosteric activation, a phenomenon commonly witnessed in many enzymes, including proteases^37^. We also examined the property of the substrate glutamate (Glu). Given the relatively low density in the GLA region of both FIX and FX in our cryo-EM maps (**Fig. 6a, b**), and considering that there are 13 Glu residues within the FIX GLA region^38^, we postulate that the observed density might represent an average of the densities of Glu residues located at various positions. Based on our prior knowledge and supported by the AlphaFold3 structure (data not shown), we primarily constructed Glu53, Glu54, and several adjacent residues in our model.

A previously reported glutamate-binding motif (GLYGYSWDMMVH_393-404_)^39^ is located at the linker region between TMD and PBD, suggesting that multiple interactions within the PBD can influence the stability of the luminal domain with TMD, consequently, affect substrate binding and enzyme activity (**Fig. 2f–h**). Additionally, the positions of various pathological mutations, such as W501S, Q503R, G558R, and T591K^20^, are located at the interface between PBD-1 and PBD-2 (Supplementary Fig. 9). This discovery indicates that these mutations probably disrupt conformational stabilities without directly impairing the propeptide release as previously reported, thereby reinforcing the importance of the coupling mechanism in regulating GGCX function. During the revision of our manuscript, two independent groups reported the apo structure of GGCX^40, 41^. Unfortunately, our efforts to determine a high-resolution cryo-EM structure of apo GGCX were unsuccessful, probably due to significant conformational flexibility in the luminal domain (Supplementary Fig. 10b). Superimposition of GGCX^WT^•FIX with the reported apo structures revealed that the PDB-2 undergoes a ∼20° rotation away from the active center in the apo state, disrupting the interactions between H2 and Cap (Supplementary Fig. 10c). This suggests that in the unbound state, the PBD-2 adopts a more open conformation, primed for binding with propeptide of VKDP. Upon binding, PBD-2 moves towards the active center and forms hydrogen bonds with Cap. Following vitamin K binding, the ternary complex assembles, initiating the carboxylation of substrate. Thus, the conformational transduction between H2 and Cap appears to be a VKDP-binding-induced structural rearrangement, which is crucial for facilitating the carboxylation reaction.

In the context of vitamin K, the oxidized vitamin K (VKO) is converted back into vitamin K hydroquinone (VKH_2_) by vitamin K epoxide reductase (VKOR) and vitamin K reductase (VKR) through a two-step reduction process^42^, thus completing the vitamin K cycle^43^. Mutation such as F299S and S300F were discovered in family with pseudoxanthoma elasticum-like phenotypes^44^, and genotype-phenotype examination proved that S300 and F299 are essential for vitamin K epoxidation^20^. Our structures have first demonstrated that these residues are involved in the naphthoquinone coordination (**Fig.4c**). Besides, we also conclude that the isoprenyl side-chain functions to bind the vitamers to the TMD within the ER membrane, but not participate in oxygenation chemistry^15^. The two main forms of natural vitamin K, phylloquinone and menaquinones, are variably distributed in mammalian tissues^45^. Given that MK-4 is notably abundant in kidney^46^ and our proteins were purified from HEK293F cells, we considered the endogenously purified vitamin K molecule in our complex to be mainly MK-4 but could not exclude other possibilities. Recent research has emphasized the significant role of vitamin K as a powerful anti-ferroptotic agent^47^, emphasizing the role of vitamin K in cellular processes and its potential applications in various fields.

Finally, the coupling between carboxylation and epoxidation activities constitutes a critical aspect of this study. In the absence of the substrate glutamate, VKH epoxidation does not take place; however, in *Leptospira* homologs where Lys217 is not conserved, the “substrate activation” mechanism does not occur, indicating that Lys217 is the key residue for the coupling mechanism. Our identification of the Cap suggests that Lys217 acts to mediate the conformational movement of vitamin K-binding to PBD via Cap-H2 transduction mechanism (**Fig. 5c–f**), rather than modulating the pKa of Lys218 as previously reported^48^. Meanwhile, His160 has dual functions in GGCX activity: on one hand, its imidazole side chain participates in the formation of the pocket for vitamin K menaquinone (**Fig. 4c**); on the other hand, it is directly hydrogen bonded with the γ-carboxyl group of FIX-Glu53 (**Fig. 5b**). Hence, His160 is essential for mediating the coupling of the two reactions. Our findings suggest that deprotonation of VKH would facilitate the deprotonation of glutamate via these two main strategies, highlighting the complex regulation of these enzymes.

In conclusion, our study provides novel insights into the molecular mechanisms of vitamin K-dependent carboxylation, spotlighting key regulatory elements such as propeptide binding and coupling of reactions. A deeper comprehension of these processes is of utmost importance for elucidating the physiological significance of VKDP carboxylation, which underpins both anticoagulant therapy and broader aspects of cellular regulation. Further studies will be necessary to illustrate whether the mechanisms governing glutamate residue carboxylation are distributive or processive, the auto-carboxylation ability of GGCX^49^, and how oxidated VKO is released from GGCX. Importantly, the findings presented here open up new avenues for further exploration into the intricate mechanisms that govern vitamin K metabolism and its role in health and disease.

## Materials and Methods

### Plasmid construction and transient expression

The optimized cDNA sequences encoding the full-length human GGCX, FIX, and FX were subcloned into the pCDNA3.1(+) vector with a C-terminal His-Flag tag. All the point mutations were generated by a standard two-step PCR and homologous recombination. Overexpression of GGCX or GGCX-substrate complexes was carried out in HEK293F cells (Invitrogen) cultured in SMM 293T-II medium (Sino Biological Inc.) at 37°C under 5% CO_2_ with 130 rpm in a Multitron-Pro shaker (Infors). When the cell density reached 2.0 × 10^6^ cells/mL, 2 mg of the total plasmids with 4mg of polyethylenimine hydrochloride (MW40,000, Polysciences) was transiently transfected into each liter cells. After 48 hours, the cell pellets were harvested by centrifugation at 2000 rpm for 20 min.

### Recombinant protein purification

For purification of GGCX, GGCX^AA^•FIX, and GGCX^AA^•FX complexes, the collected cell pellets were thoroughly resuspended in lysis buffer (20 mM HEPES-K pH 7.4, 150 mM NaCl, 10% Glycerol, 1 mM MgCl_2_, 1 mM EDTA). Subsequently, 1% GDN detergent, protease-phosphate inhibitors, and DNase were added and mixed homogeneously. The mixture was incubated at 4□ for approximately two hours and then centrifuged at 30,000 rpm for an hour. Next, ANTI-FLAG^®^ M2 Affinity Gel beads were equilibrated with the lysis buffer containing 0.01% GDN, and the supernatant obtained after ultra-centrifugation was loaded to the equilibrated FLAG beads and incubated at 4 for an hour. The beads were washed three times with lysis buffer supplemented with 0.01% GDN, 0.1mM PMSF, and 1 mM ATP. The bound protein was eluted with lysis buffer supplemented with 0.2 mg/mL Flag-peptide and concentrated to approximately 2 mL. The eluted protein was applied to size exclusion chromatography (Superose-6 Increase 10/300 GL, Cytiva) in the lysis buffer containing 0.01% GDN.

### Cryo-EM sample preparation and data acquisition

Holy-carbon gold grids (Quantifoil Au 400 mesh, R1.2/1.3) were glow-discharged in either Solarus 950 plasma cleaner (Gatan) or PELCO easiGlow (Ted Pella) before cryo-EM sample preparation. 4 μL aliquots of freshly prepared GGCX^WT^•FIX, GGCX^AA^•FIX, and GGCX^AA^•FX complexes (4 mg/mL) were applied on the glow-discharged grids, blotted with filter paper (Whatman No.1) with force set to –2 for 0.5 s at 4°C and 100% humidity, and plunge-frozen in the liquid ethane using a Vitrobot Mark IV (Thermo Fisher Scientific).

For GGCX^WT^•FIX complex, the cryo-grids were screened on a 200 kV Talos Arctica equipped with an FEI Ceta detector. Data collection was carried out using Titan Krios G3 (FEI) with a K3 Summit (Gatan) operating at 300 kV. Images were recorded in the super-resolution mode at a nominal magnification of 81,000× and a dose rate of 15□e^−^/s/pixel. A GIF BioQuantum energy filter (Gatan), with a slit width of 20□ eV, was used at the end of the detector. The defocus range was set from –0.8 to –1.2□ μm. The total exposure time was 4.58 s, and intermediate frames were recorded every 0.14 s. A total of 32 frames per image were acquired.

For GGCX^AA^•FIX and GGCX^AA^•FX complexes, the cryo-grids were screened on a 200 kV Glacios^TM^ 2 equipped with Falcon 4i (Thermo Fisher Scientific). Data collection was carried out using Krios G4 (Thermo Fisher Scientific) operating at 300 kV. Images were recorded with a Gatan K3 Summit detector in the super-resolution mode at a nominal magnification of 105,000× with a calibrated pixel size of 0.85 Å and a dose rate of 17e^−^/s/pixel. A GIF BioContinuum filter (Gatan), with a slit width of 20 eV, was used at the end of the detector. The defocus range was set from −0.8 to −1.8 μm. The total exposure time was 2.00 s, and intermediate frames were recorded every 0.06 s. A total of 32 frames per image were acquired. All movies were recorded semi-automatically using the EPU software. The statistics for data collection are summarized in Extended Table S1.

### Imaging processing

A total of 4598, 13529, and 6911 movie stacks were recorded for GGCX^WT^•FIX, GGCX^AA^•FIX, and GGCX^AA^•FX complexes, respectively. We processed the cryo-EM data using the CryoSPARC^50^. Motion correction of the micrographs and the contrast transfer function (CTF) parameters were finished using patch motion correction and patch CTF estimation, respectively. Blob-picking, 2D classification, Ab-initio reconstruction, heterogeneous refinement, homogeneous refinement, non-uniform refinement, and 3D classification were sequentially done according to the CryoSPARC data procession protocol. The resolution of the map was estimated by Fourier shell correlation (FSC) analysis at a correlation cutoff value of 0.143. Local resolution maps were analyzed using Local Resolution Estimation, resulting in the resolutions of 2.78, 2.59, and 2.58 Å for GGCX^WT^•FIX, GGCX^AA^•FIX, and GGCX^AA^•FX complexes, respectively. Workflow of the data processes were illustrated in the Supplementary Figs. 2 and 8.

### Model building and refinement

The initial model of GGCX was built by AlphaFold2^51^. The initial model was then fitted into the cryo-EM map using ChimeraX^52^. FIX and FX were manually added into the initial models and adjusted manually using Coot^53^. Vitamin K-restraint files for refinement were generated by phenix.elbow^54^. The model refinements of all three datasets against the corresponding maps were performed using PHENIX in real space with secondary structure and geometry restraints^55^. The final structures were validated through examination of the Clash scores, Molprobity scores and statistics of the Ramachandran plots by PHENIX^56^.

### Simulation system design and configuration

We established three systems: GGCX-MK-4, truncated GGCX (GGCXΔArch)-MK-4, and GGCXΔArch-FIX-MK-4. The cryo-EM model of WT-GGCX was preprocessed for MD simulation with deprotonation of lysine residues Lys217 and Lys218. Molecular docking of MK-4 respect to the modified GGCX was carried out using AutoDock Vina^57^ to generate the initial structures of the protein-ligand complexes. To avoid an artificial interaction between termini, the truncated structure was capped with an acetyl (ACE) group at the N-terminus and an N-methylamide (NME) group at the C-terminus. To mimic the physiological environment of GGCX, the complexes were embedded into a lipid bilayer composed of POPC mixed with a minor proportion of cholesterol using the CHARMM-GUI^58^. After solvation and neutralization with 150 mM KCl, the total number of atoms in each system reached approximately 260,000.

The force fields leaprc.protein.ff19SB^59^, leaprc.lipid21^60^, leaprc.water.tip3p^61^, and the Generalized Amber Force Field (leaprc.gaff2)^62^ were employed to parameterize the protein, lipids, water molecules (including ions), and MK4, respectively. Force field parameters for MK4 were generated using Antechamber^63^, and the topology and coordinate files for the entire simulation system were prepared with LEap^63^. In order to accelerate simulation speed, SHAKE and hydrogen mass repartitioning^64^ were used to allow the implementation of a 4 fs timestep.

### Simulation process

Following energy minimization to relax steric clashes, a 4 ns pre-equilibration was run in the NPT ensemble to naturalize the lipid bilayer with the protein and ligand fixed by a force k = 100 kcal/mol/Å^2^. Then restraints were gradually removed to allow the system to approach a low-energy state. Under periodic boundary conditions (PBC), all pre-equilibrations were performed at 300 K (1 atm) and the van der Waals cutoff of 10 Å, using the Particle Mesh Ewald (PME) method to calculate electrostatic interactions. Tiny fluctuations of the system volume indicated that membrane and solvent molecules were well equilibrated, and thus we fixed volume to for the subsequent production simulations for each system. The minimization, pre-equilibration and equilibrium simulation were performed with OpenMM^65^.

Considering that the timescales of conformational changes relevant to ligand unbinding process in the GGCXΔArch -FIX-mk4 system exceed the limit of conventional MD (cMD) simulations, we conducted accelerated MD (aMD)^66^ simulations to sample the conformational transition in this system. By adding a boost potential ΔV(r) to the total potential and dihedral potential, aMD encourages the protein to evolve faster without altering the shape of potential energy surface:

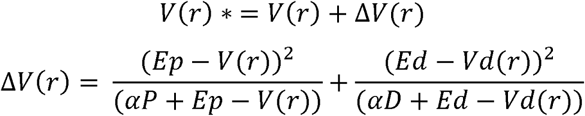

Where *V(r)* and *Vd(r)* are the normal total and torsion potential respectively. Ep and Ed are average potential and dihedral energies that serve as reference threshold values. The terms αP and αD denotes factors that determine inversely the strength of the boost that is applied. After testing some combinations of parameters, we found that the total number of atoms scaled by 0.20 seemed to work well without perturbing the system stability. Subsequently, we extended the aMD simulations to ∼160 ns in three repeats using the binary file “pmemd.cuda” in Amber23.

### Evaluating motions of Cap and H2

By analyzing per-residue RMSF and functional domain segmentation, we identified three regions: 1) the TMD with small RMSFs, which was used as a spatial reference to eliminate translational and rotational degrees of freedom from the trajectories; 2) the flexible Cap (residues 237-254) which exhibits dynamics relative to the ligand, and 3) the second α-helix in PBD-2 (H2, residues 510-524) whose motion is associated with the Cap dynamics. To evaluate the opening motion of the Cap, the center of mass of the Cap region was computed and compared to its initial position, yielding a metric *d_r_*. For H2, principal component analysis (PCA) was applied on the coordinates of Cα atoms in each frame, with the first principal component (PC1) representing the dominant direction of H2. The angle between the PC1 vector of the trajectories and that of the initial structure (cos *θ*) was measured to quantify H2 movement. We used MDTraj^67^ for reading, superposing and RMSD calculation of MD trajectories. The RMSFs was computed based on the “rmsf” command in CPPTRAJ^68^, and PCA was implemented with Scikit-learn^69^.

### Substrate purification

The cDNAs for coagulation factor IX (FIX) and other substrates (FVII, FX, PS, GAS6) were subcloned into the pET-21c(+) vector with a N-terminal maltose-binding protein (MBP)-His tag. The expression was induced by IPTG in *E.coli*. Purification was performed using Ni Smart Beads 6FF (Smart-Lifesciences). After a 15-min incubation of the supernatant with the beads, the samples were washed and eluted with 250 mM imidazole. The eluted protein was concentrated with Amicon® Ultra-15 ultrafiltration tube (Millpore), followed by ion exchange chromatography (SOURCE-15Q) with a gradient of NaCl (50 mM–1 M) in 20 mM Tris-Na, pH 8.0. Fractions containing target proteins were collected, concentrated, and applied to gel filtration chromatography (Superdex-200 10/300 GL, Cytiva) in the lysis buffer. The peak fractions were concentrated and stored at −80□.

### *In vitro* carboxylation reaction assay

A western blotting-dependent *in vitro* detection system for GGCX carboxylase activity was established. 2.5 μg of GGCX and its substrate (containing the propeptide and GLA region) were mixed in a molar ratio of 1:30 and incubated on ice for an hour. MK-4 was reduced to VKH_2_ with DTT reduction. 250 μM of VKH_2_ and 920 μM of NaHCO_3_ were added to the reaction system, which was adjusted to a total volume of 50 μL with lysis buffer with 0.01% GDN. The reaction was carried out at 20°C for an hour, and the reaction mixtures were loaded onto electrophoresis and visualized by western blotting using Gla antibody (Cat#3570, BioMedica Diagnostics).

### Cell-based carboxylation activity assay

HEK 293T cells are maintained in DMEM supplemented with 10 % FBS and 1% penicillin-streptomycin at 37 °C with 5 % CO_2_. Cells were seeded in a 12-well plate at 70% confluency before transfection. Transient transfection of GGCX and its full-length substrates at a molar ratio of 3:5 was accomplished using Lipofectamine 3000 Reagent (L3000015, Thermo Fisher) with 10 µM vitamin K. After 48 hours of incubation, cells were lysed in lysis buffer containing 1% GDN at 4 for an hour. The supernatant was collected by centrifugation and analyzed via Western Blotting using GGCX antibody (16209-1-AP, Proteintech) and Gla antibody (Cat#3570, BioMedica Diagnostics), respectively.

### Pull-down assay

Pull-down assays were carried out using Flag-immobilized GGCX in combination with either GFP- or MBP-fused FIXQ/S region of FIX (aa29–92). For pull-down assays of FIX mutants (WT, F31G, H34R, and L41W), the substrates were fused with super-folded GFP (sfGFP) at the N-terminus and purified as previously described. FLAG beads were used to capture 2.5 μg Flag-tagged GGCX through a 30-minute incubation on ice. Subsequently, an excess amount of sfGFP-fused FIX proteins was introduced and further incubated for an additional 30 minutes. The samples were eluted with 0.2mg/mL flag-peptide and subsequently subjected to immunoblot analysis for the western-blot detection of GGCX and fluorescent detection of sfGFP. For pull-down assays of GGCX variants (WT, K217A, K218A, K217A & K218A), all the procedures remained identical, with the exception that MBP-fused FIX was employed and analyzed via western blotting using MBP antibody (E022240, EARTHOX).

### Western blotting

The samples of GGCX carboxylation reaction mixture were resolved by SDS-PAGE and transferred to Polyvinylidene fluoride (PVDF) membranes (IPVH00010, Millipore). The membranes were blocked by 5% (w/v) Skim Milk (232100, BD) at room temperature for one hour and incubated with the primary antibodies as indicated above at 4°C overnight with an appropriate dilution ratio. Then the membranes were washed with the TBST buffer (25 mM Tris, pH 8.0, 150 mM NaCl, and 0.05% (w/v) Tween-20) for three times. The Horseradish Peroxidase (HRP) conjugated-secondary antibodies (P03S02M or P03S01M, Gene-Protein Link) were incubated for 1 hour and washed similarly, and visualized under automatic chemiluminescence imaging analysis system (Tanon).

### Surface plasmon resonance

The equilibrium dissociation constant (*K_D_*) values for GGCX-FIX and GGCX-vitamin K interactions were determined using Surface Plasmon Resonance (SPR) experiments conducted on Biacore 8K+ instrument (Cytiva). Recombinant GGCX proteins were diluted in sodium acetate solution (pH 5.5) with 0.01% GDN to a final concentration of 10 ng/μL. A total volume of 100 μL GGCX proteins was immobilized on CM5 sensor chips via amine coupling. For GGCX-FIX (WT, F31G, L41W, and H34R) interactions, FIXQ/S analytes within the concentration range of 78 nM to 5 μM were injected at a rate of 30 μL/min for 120 s in single-cycle mode. During the dissociation phase, the running buffer (20 mM HEPES-K pH 7.4, 150 mM NaCl, 1 mM MgCl_2_, 1 mM EDTA) was injected at a flow rate of 30 μL/min for 240 s. For GGCX-vitamin K interaction, DMSO-dissolved vitamin K (0.78 μM–400 μM) was injected at a rate of 30 μL/min for 60 s in single-cycle mode, followed by the injection of the running buffer with 5% DMSO at a flow rate of 30 μL/min for another 60 s. All data were recorded at 25 °C and analyzed using the Biacore 8K+ Evaluation Software.

## Data availability

The cryo-EM maps of GGCX•FIX, GGCX^AA^•FIX, and GGCX^AA^•FX have been deposited at the EMDB with the accession codes of EMD-62862, EMD-62863, and EMD-62864, respectively. The corresponding atomic models have been deposited at the RCSB PDB with the accession codes of 9L6Q, 9L6R, and 9L6S, respectively.

## Author contributions

This work was performed by all authors. J.H., J.Q., and L.T. conceived and designed the experiments. Q.Z., J.X., Y.M., and K.P. performed the protein purification and the majority of biochemical experiments. D.C., J.H., Q.W., and G.W. performed the cryo-EM sample preparation and data analysis. Y.L. and Q.Y. performed the computational experiments. J.H., J.X., and Q.Z. wrote and edited the manuscript with input from all authors.

## Acknowledgements

Cryo-EM data were collected either at Peking University Health Science Center Cryo-Electron Microscopy Facility or the Cryo-EM platform of Peking University. This work was supported by grants from the National Key R&D Program of China (2022YFC2702904, 2023YFC2705902 and 2022YFC2703800), the National Natural Science Foundation of China (32470880 and 32401042), Key Clinical Projects of Peking University Third Hospital (BYSYZD2023033), and China Postdoctoral Science Foundation (2023M730126).

