## Supplemental Information for "Molecular basis of vitamin K-dependent protein γ-glutamyl carboxylation"

**Supplementary Fig. 1–10 and legends**

**Supplementary Tables 1–2**

**Supplementary Video S1–2 titles**

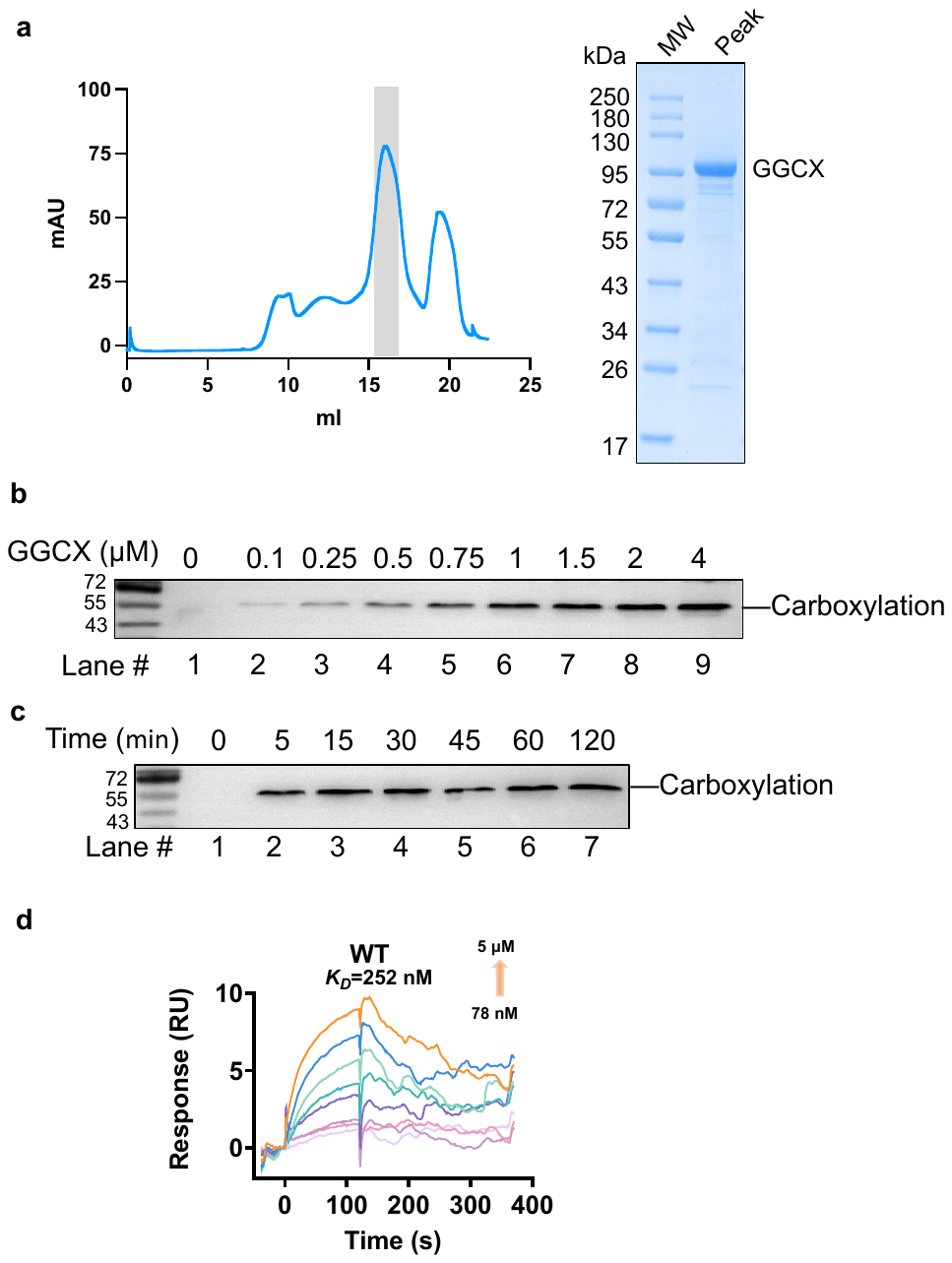

**Supplementary Fig. 1 | Purification and characterization of GGCX. a,** Size-exclusion chromatography (left) and the Coomassie blue staining of SDS-PAGE (right). The gray-shaded fractions were used for activity analysis. **b–c,** Concentration-dependent (**b**) and time-dependent (**c**) in vitro reaction using Factor IX (FIX) as the substrate. **d,** Real-time response of FIX at different concentrations against GGCX by SPR.

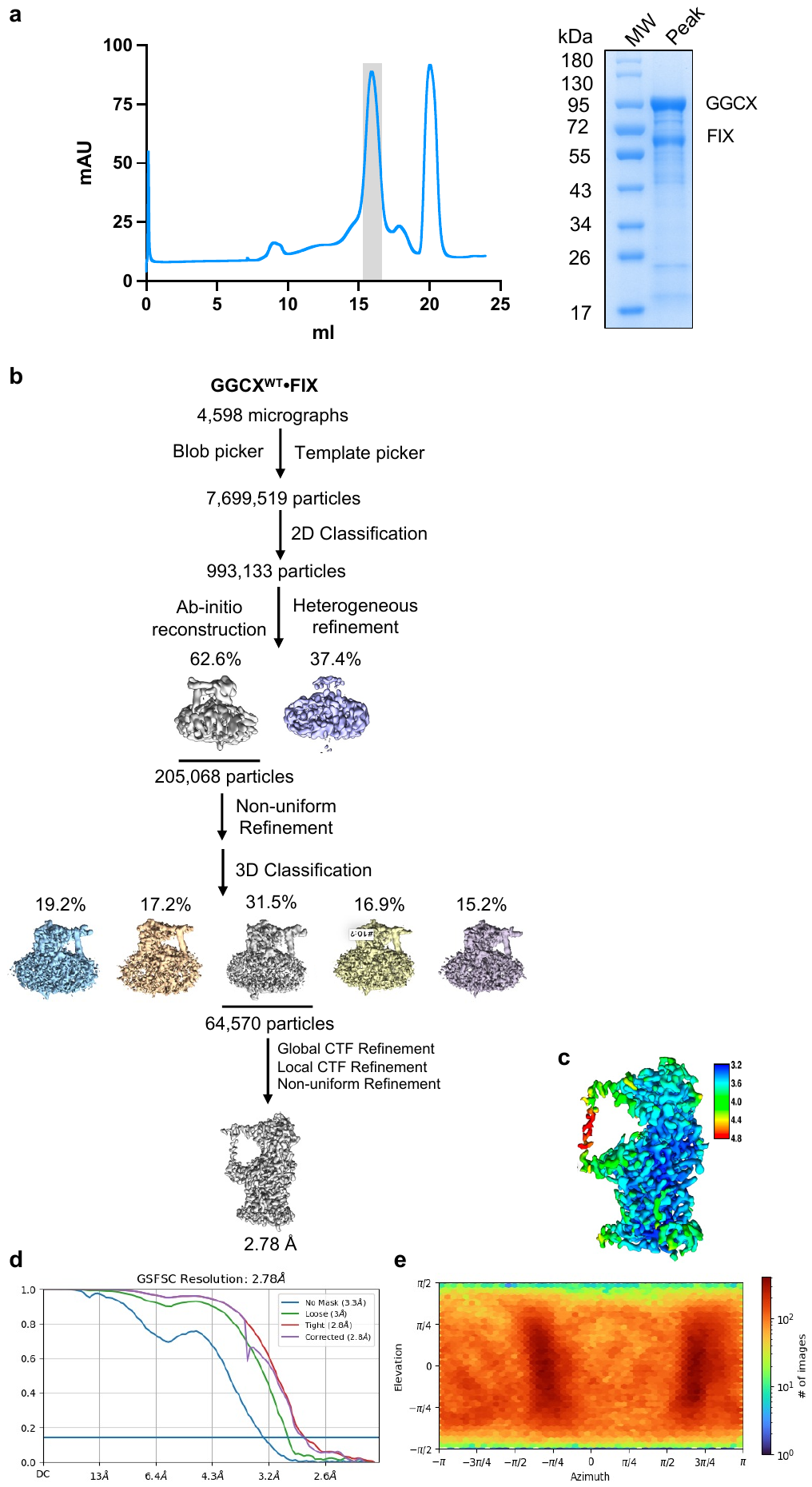

**Supplementary Fig. 2 | Cryo-EM structural analysis of GGCX•FIX complex. a,** The purification of GGCX•FIX complex. Size-exclusion chromatography (left) and the Coomassie blue staining of SDS-PAGE (right). The gray-shaded fractions were used for cryo-EM analysis. **b,** Cryo-EM data processing procedure accomplished in CryoSPARC. The representative results of the 2D classification were shown. **c,** Local resolution map of the 2.78-Å 3D map. **d,** Gold-standard Fourier shell correlation (FSC) of two independent half 3D maps of GGCX•FIX•vitamin K complex. **e,** Angular distribution of raw particles used in CryoSPARC 3D reconstruction.

**Supplementary Fig. 3 | Representative cryo-EM densities for GGCX•FIX•vitamin K complex. a–c,** Density map and model in selected regions of GGCX (a), substrate FIX (b), and the glycosylation sites (c). The sidechains of some residues are shown as sticks.

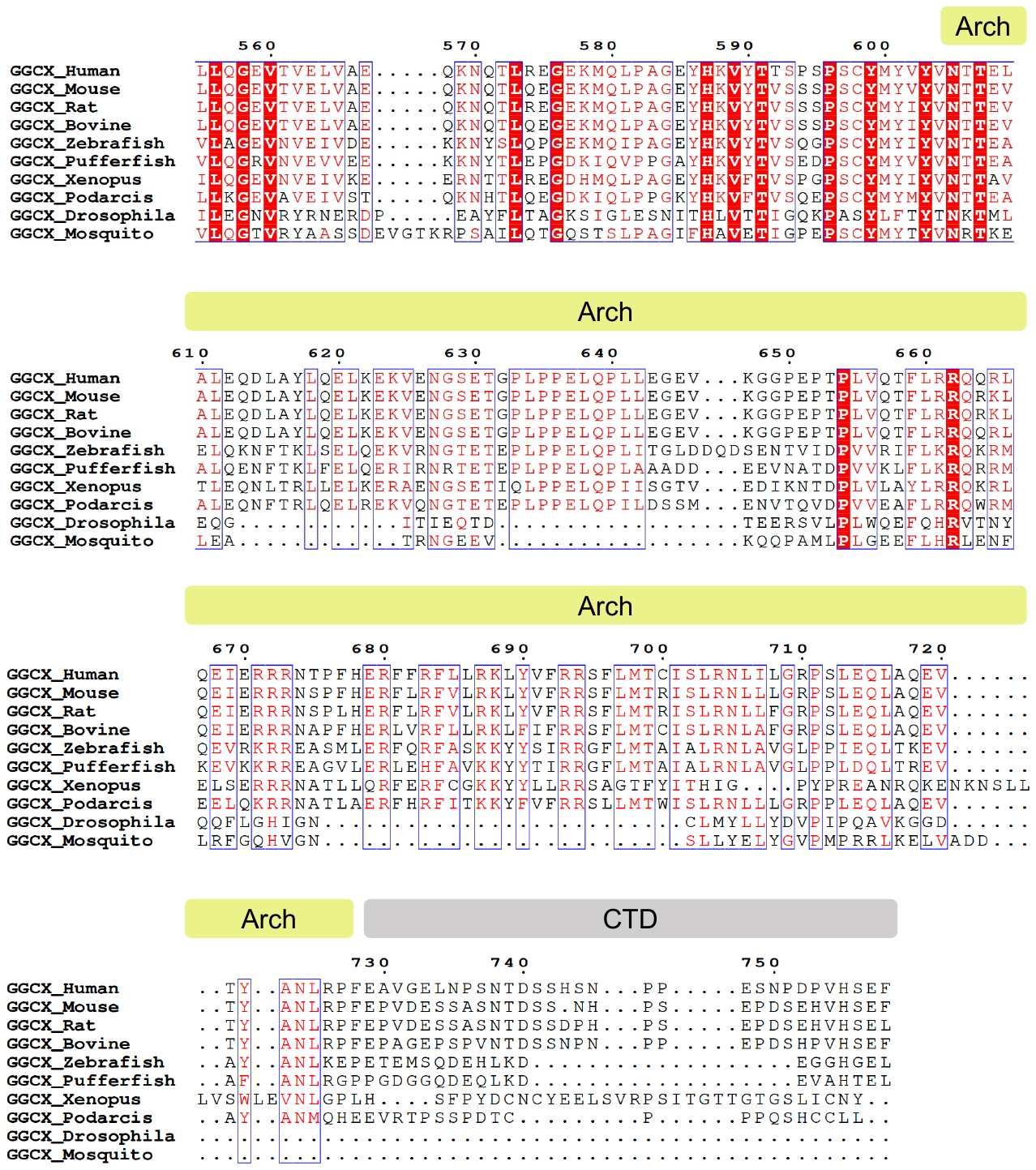

**Supplementary Fig. 4 | Sequence alignment of GGCX homologs.** Alignment of amino acid sequences of GGCX homologs generated with Clustal Omega (Uniprot identifiers: P38435, Q9QYC7, O88496, Q07175, A0A8M1RIV5, H3CUP2, A0A6I8PZ28, A0A670IZL4, Q9NDA0, Q7Q5L8). The conserved Arch domain and the unconservative C-terminal domain (CTD) are shown.

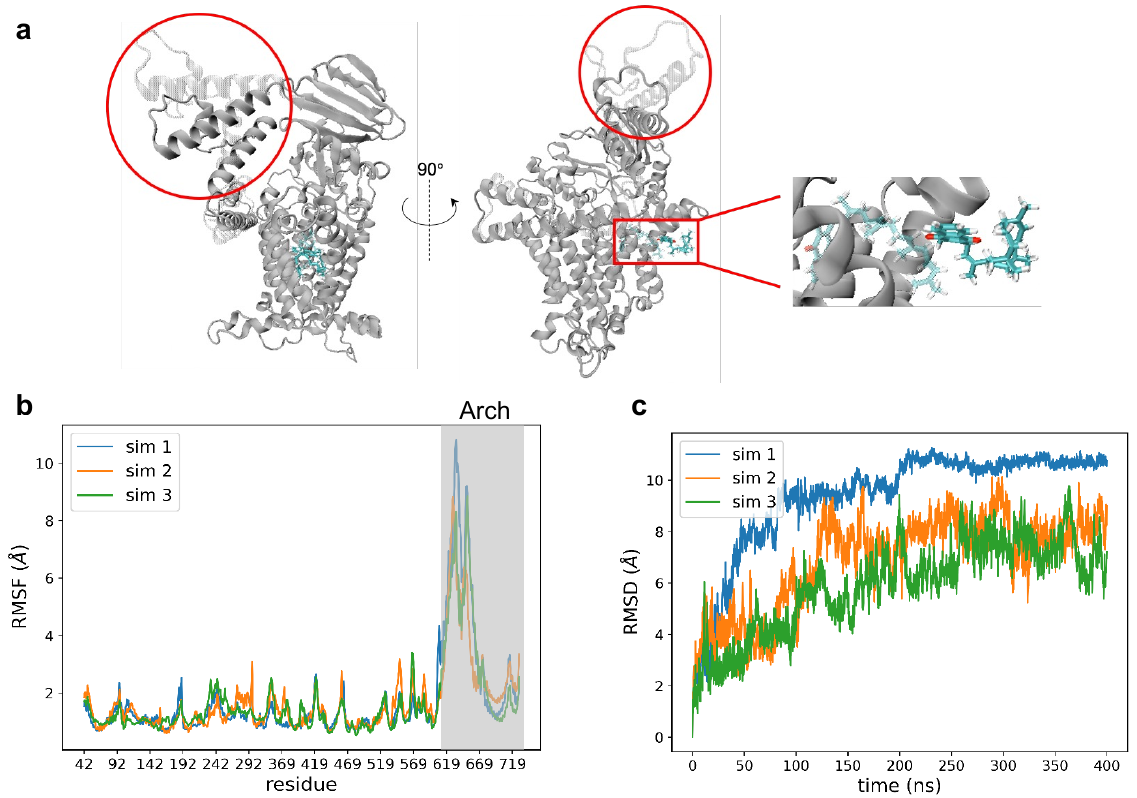

**Supplementary Fig. 5 | The flexibility of the ER luminal domain in apo structure. a,** Structural representation of the GGCX-MK4 system. The left and middle panels display side views of the final state from the MD simulations, with the initial configuration of the Arch region shown in transparent mode. Conformational changes are highlighted with red circles. The right panel shows the conformational change of binding Vitamin K (colored blue) with initial conformation shown in transparent. **b,** Per-residue root-mean-square-fluctuation (RMSF) of GGCX from the MD simulations without the C-terminal residues (ΔC). The Arch domain is indicated in light gray. **c,** Root-mean-square-deviation (RMSD) of the Arch region during 400-ns MD simulations. Results from three independent simulation replicates are shown in blue, orange, and green, respectively.

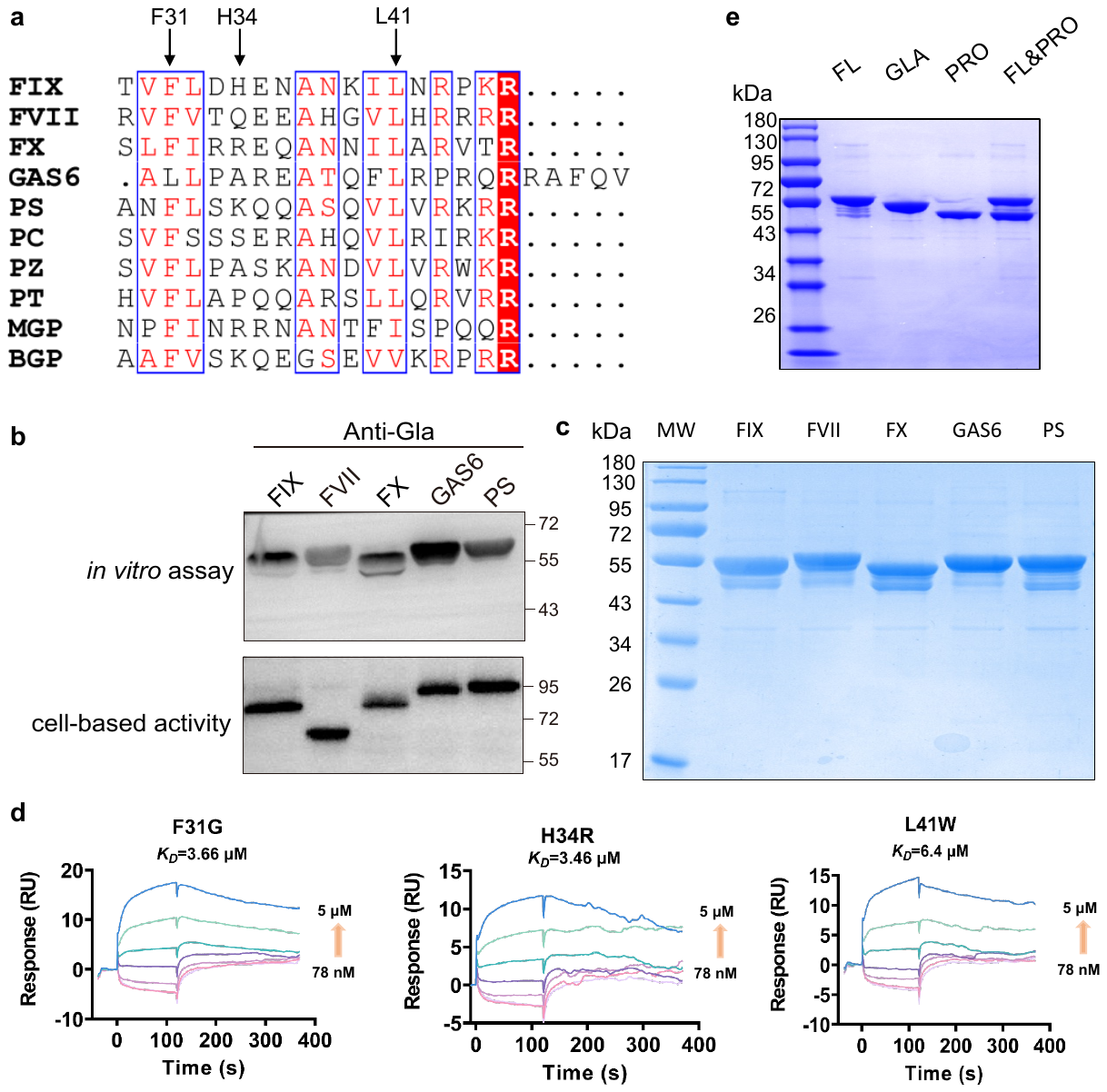

**Supplementary Fig. 6 | The binding and carboxylation of various substrates. a,** Sequence alignment for various substrates of GGCX. The conserved residues of substrates are labeled. **b–c,** Carboxylation activity of different substrates by *in vitro* reaction (b) under normalized input substrates (c). FIX, coagulation factor IX; FVII, coagulation factor VII; FX, coagulation factor X; GAS6, growth arrest-specific protein 6; PS, protein S. **d,** Real-time response of the mutant substrates (F31G, H34R, and L41W) at different ligand concentrations against GGCX by SPR. **e,** Inputs of substrates were normalized for the *in vitro* reaction in Fig. 3g.

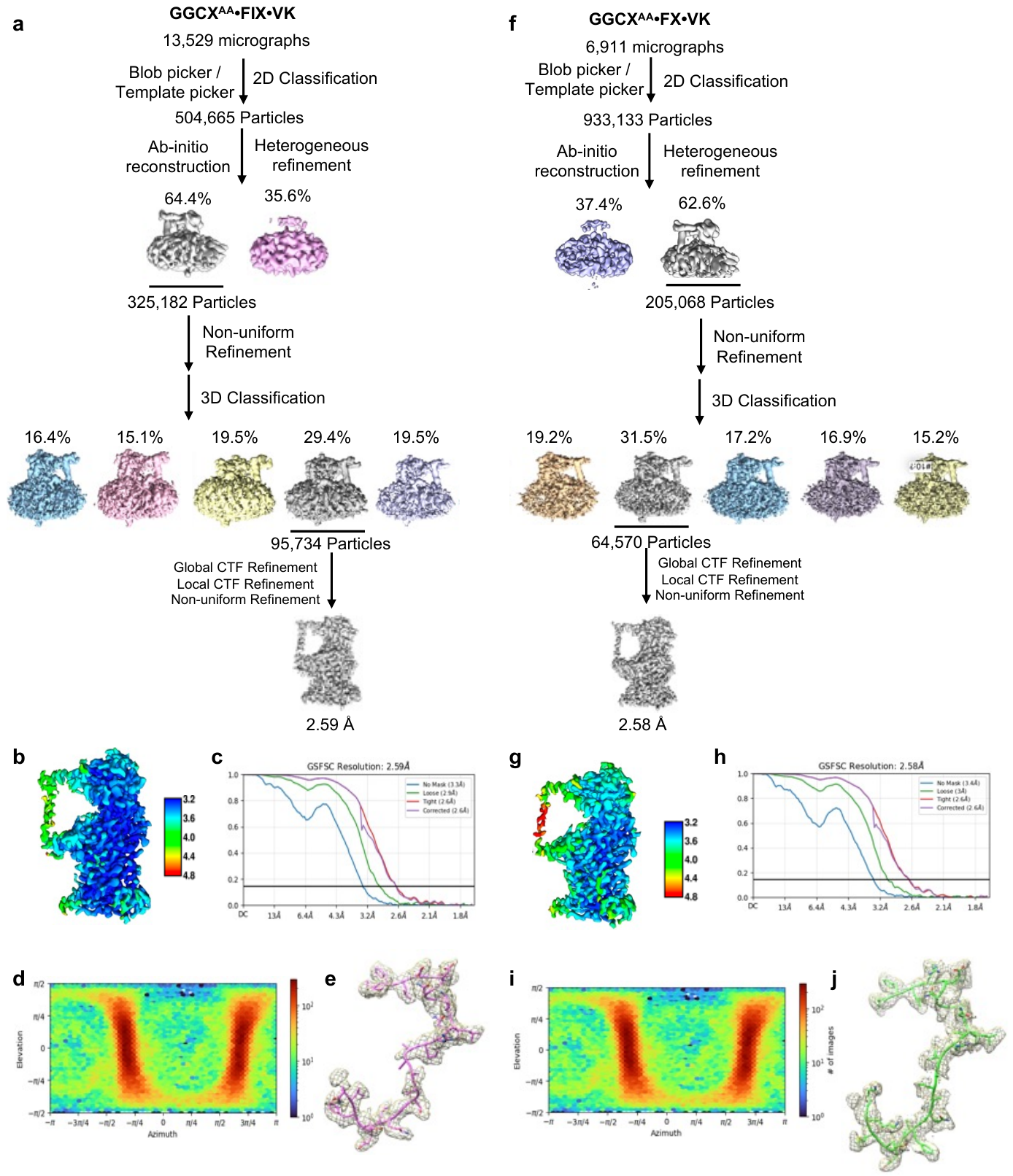

**Supplementary Fig. 7 |** **Cryo-EM structural determination of FIX- and FX-bound mutated GGCX. a,** Cryo-EM data processing procedures for GGCX-K217A&K218A (GGCX^AA^) in complex with FIX and MKH_2_-4. **b,** Local resolution map of the GGCX^AA^•FIX•MKH_2_-4 complex at 2.59-Å. **c,** Gold-standard Fourier shell correlation (FSC) of 3D map of the complex. **d,** Angular distribution of raw particles used in CryoSPARC 3D reconstruction. **e,** EM density map and the corresponding model for the substrate FIX. **f–j,** Similar procedures were applied for structural determination of GGCX^AA^•FX•MKH_2_-4 complex at 2.58-Å.

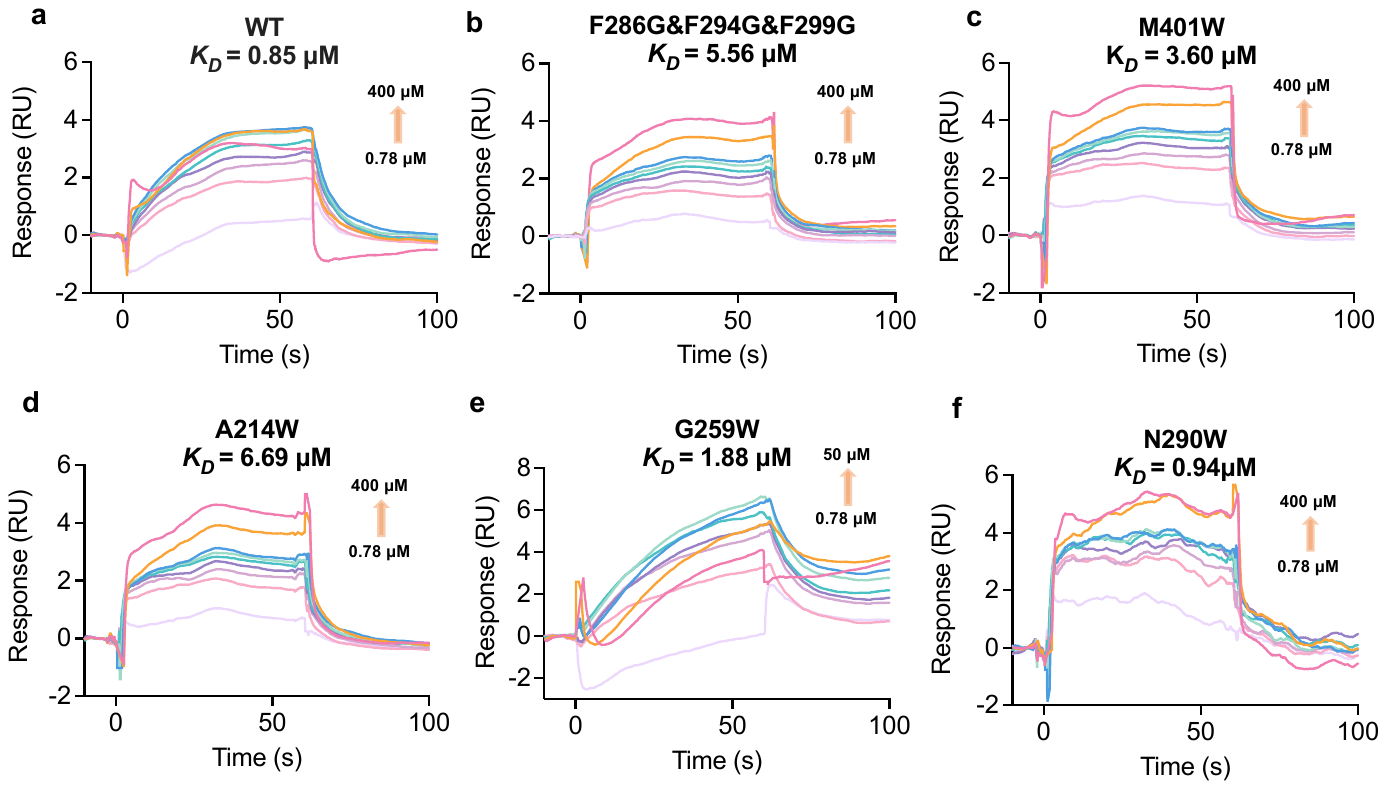

**Supplementary Fig. 8 | SPR analysis of vitamin K binding to GGCX.** Real-time response of vitamin K at different ligand concentrations against the mutant GGCX Concentrations used are shown. KD values are labelled

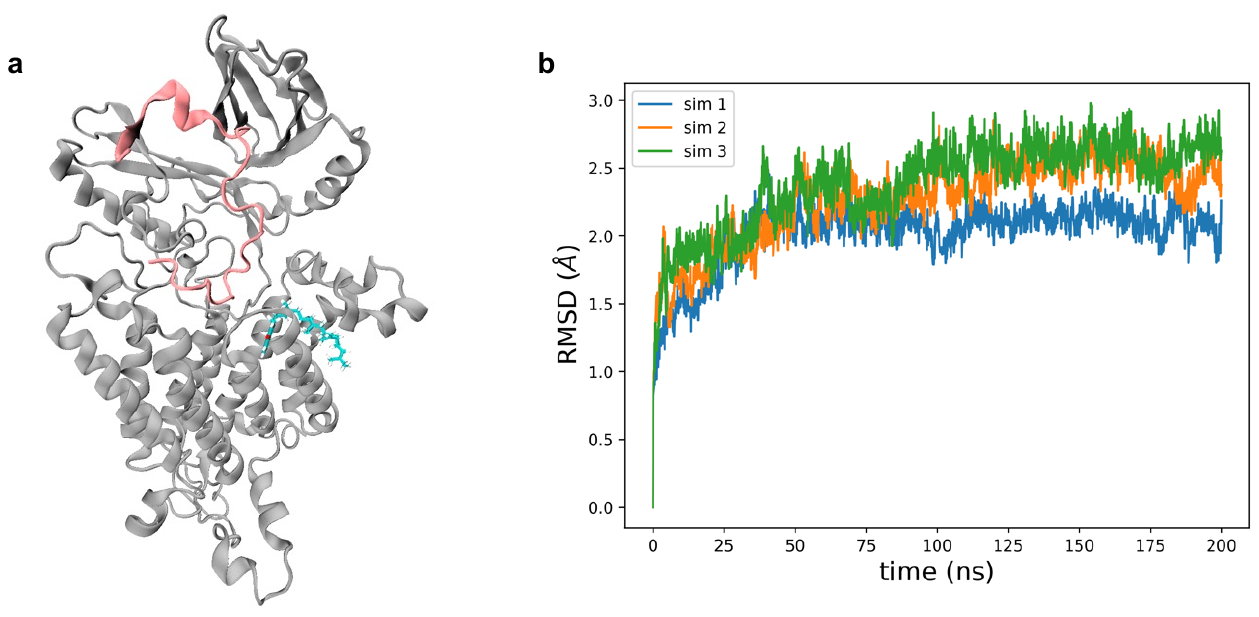

**Supplementary Fig. 9 | Conventional MD simulation** **using a GGCXΔArch-FIX-MK-4 system. a,** The structural illustration of GGCX∆Arch-FIX-MK-4 system with FIX colored in pink. **b,** Time course plot for GGCX∆Arch and FIX RMSD changes during three independent repeats of 200-ns simulations.

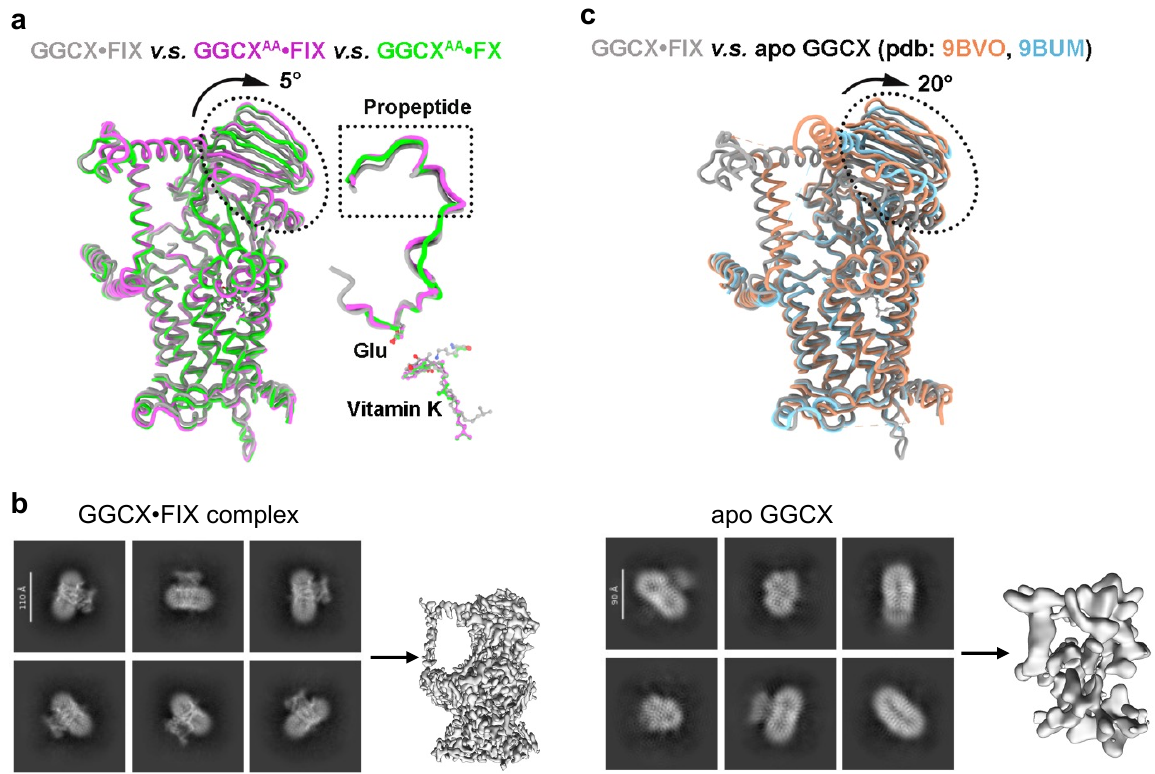

**Supplementary Fig. 10 |** **Conformational flexibility of GGCX. a,** Superimposition of GGCX^AA^•FIX and GGCX^AA^•FX with the wild-type GGCX•FIX structure. The PBD-2 domain is highlighted by a dashed ellipse. The overlay of bound substrates and vitamin K across the three structures is shown in the right panel. The propeptides of the substrates are highlighted in a dashed box. **b,** The representative results of the 2D classification and 3D map of the resolved GGCX•FIX complex (left) and that of apo GGCX (right). **c,** Superimposition of GGCX•FIX with apo structures (PDB ID: 9BVO and 9BUM). The PBD-2 domain is highlighted by a dashed ellipse.

**Supplementary Table 1. Cryo-EM data collection, refinement and validation.**

|  | **GGCX**•**FIX complex (EMD-62862; PDB ID: 9L6Q)** | **GGCX^AA^**•**FIX complex (EMD-62863; PDB ID: 9L6R)** | **GGCX^AA^**•**FX complex (EMD-62864; PDB ID: 9L6S)** |
| --- | --- | --- | --- |
| **Data collection and processing** | | | |
| Voltage (kV) | 300 | 300 | 300 |
| Microscope | FEI Titan Krios G3 | Thermo Fisher Scientific Krios G4 | Thermo Fisher Scientific Krios G4 |
| Camera | Gatan K3 Summit with energy filter | Gatan K3 Summit with energy filter | Gatan K3 Summit with energy filter |
| Magnification (calibrated) | 81,000 × | 105,000 × | 105,000 × |
| Electron exposure (e^–^/Å^2^) | 60 | 40 | 40 |
| Exposure rate | 13.10 e^–^/Å^2^/s | 23.53 e^–^/Å^2^/s | 23.53 e^–^/Å^2^/s |
| Number of frames per micrograph | 32 | 32 | 32 |
| Energy filter slit width (eV) | 20 | 20 | 20 |
| Defocus range (-μm) | 0.8-1.2 | 0.8-1.8 | 0.8-1.8 |
| Pixel size (Å) | 1.07 | 0.85 | 0.85 |
| Micrographs used | 4,598 | 13,529 | 6,911 |
| Initial particle images (no.) | 11,235,095 | 17,399,039 | 7,699,519 |
| Final particle images (no.) | 331,914 | 95,734 | 64,570 |
| Symmetry imposed | C1 | C1 | C1 |
| Map resolution (Å) | 2.78 | 2.59 | 2.58 |
| FSC threshold | 0.143 | 0.143 | 0.143 |
| **Refinement** | |  |  |
| Resolution (Å) at 0.143 FSC of masked reconstruction | 2.80 | 2.54 | 2.54 |
| Resolution (Å) at 0.5 FSC of masked reconstruction | 2.97 | 2.72 | 2.74 |
| Map sharpening B factor (Å^2^) | 78.5 | 68.3 | 66.1 |
| Model composition |  |  |  |
| Non-hydrogen atoms | 5,807 | 5,699 | 5,693 |
| Protein residues | 696 | 689 | 687 |
| Ligands | LIG: 1  BMA: 2  NAG: 7 | LIG: 1  BMA: 2  NAG: 7 | LIG: 1  BMA: 2  NAG: 7 |
| R.m.s. deviations |  |  |  |
| Bond lengths (Å) | 0.004 | 0.002 | 0.004 |
| Bond angles (°) | 0.809 | 0.607 | 0.639 |
| Validation |  |  |  |
| MolProbity score | 2.53 | 1.71 | 1.98 |
| Clashscore | 29.97 | 7.76 | 4.15 |
| Poor rotamers (%) | 1.85 | 1.73 | 4.05 |
| Ramachandran plot |  |  |  |
| Favored (%) | 94.74 | 97.50 | 96.76 |
| Allowed (%) | 5.26 | 2.50 | 3.24 |
| Disallowed (%) | 0.00 | 0.00 | 0.00 |
| C-beta outliers (%) | 0.00 | 0.00 | 0.00 |
| CaBLAM outliers (%) | 2.23 | 1.49 | 1.79 |

**Supplementary Table 2. Disease-related GGCX mutations.**

| **GGCX Domain** | **VKCFD-related GGCX mutations** |
| --- | --- |
| **Transmembrane Domain**  **(TMD)** | D31N P80L R83W/P D153G W157R G125R D153G W157R M174R R204C V255M S284P F299S S300F W315X R325Q Q374X |
| **Propeptide-Binding Domain**  **(PBD)** | R476C/H R485P W493C/S W501S I532T D534V G537A G558R T591K |
| **Arch Domain** | R704X |
| **Other Positions** | D31N L394R H404P |

**Supplementary Videos S1. Conformational changes of apo GGCX measured by MD simulation**. The structure of GGCX is shown in cartoon and colored gray.

**Supplementary Videos S2.** **Conformational changes of the** **GGCXΔArch-FIX measured by MD simulation.** The C-terminal region and Arch domain of GGCX are truncated (GGCXΔArch). The structure of GGCXΔArch is colored gray, and the bound FIX is shown in pink.
